# From Subconscious to Insight: Decoding the Incubation Process in Creative Problem Solving

**DOI:** 10.64898/2026.07.15.738837

**Authors:** Chatrin Phunruangsakao, Zenas C. Chao

## Abstract

Creative problem-solving is a fundamental aspect of human cognition, yet its underlying neural mechanisms remain incompletely understood. The incubation period, during which individuals temporarily disengage from conscious problem-solving efforts, has been proposed to facilitate insight through ongoing subconscious processing. Memory replay, defined as the sequential reactivation of previously encoded memories, has been implicated in a variety of cognitive processes, including learning, planning, and memory consolidation. Since insight is thought to arise from the reorganization and integration of existing knowledge, memory replay may provide a mechanism through which such restructuring occurs during incubation. This study investigated whether memory replay during incubation predicts subsequent insight and examined the replay dynamics associated with insight emergence. To this end, a novel visual sequential insight task (VSIT) was developed, and electroencephalography (EEG) data were recorded during the incubation period from 58 participants. Using a localizer model trained on EEG data acquired in a separate localization session, spontaneous memory reactivation events during incubation were decoded and subsequently used to reconstruct replay. The results demonstrate that specific patterns of memory replay are associated with successful insight, suggesting that the subconscious reprocessing of task-relevant information contributes to creative problem-solving. Furthermore, insight-related memory replay was accompanied by enhanced low-gamma-band (30–50 Hz) functional connectivity across networks involved in visual processing, memory, reward, and sensorimotor functions. Together, these findings provide evidence for the role of memory replay in supporting insight and advance current understanding of the neural mechanisms underlying creativity.

## 1 Introduction

Creativity is a multifaceted cognitive process that enables individuals to generate novel ideas or solutions [1]. A critical component of this process is insight, which refers to the sudden realization of a solution to a previously unsolved problem. Insight often occurs after a period of incubation, during which individuals disengage from conscious problem-solving efforts and allow subconscious processes to operate [2–4]. Behavioral evidence demonstrates that incubation can enhance creative problem-solving, particularly when the offline period involves rest, sleep, or engagement in an undemanding task [3–6]. These findings suggest that problem-related neural processing continues during incubation and may facilitate the emergence of novel associations and alternative solution strategies.

Several theoretical frameworks have been proposed to explain the incubation effect, including attentional, emotional, and memory-based accounts. Attentional theories suggest that disengagement from a problem reduces fixation on ineffective solution strategies, thereby increasing access to alternative approaches [2]. Emotional accounts propose that relaxation or positive affect broadens associative thinking and promotes creative cognition [7]. Memory-based accounts, in contrast, emphasize the reactivation, reorganization, and recombination of previously encoded information during offline periods. Although behavioral evidence supports each of these perspectives, the neural mechanisms through which incubation facilitates insight remain poorly understood. Among these accounts, memory-based frameworks are particularly compelling because they provide a plausible computational link between the restructuring of existing knowledge and the emergence of novel solutions [6, 8, 9].

Memory replay, defined as the sequential reactivation of neural representations associated with prior experiences, has been implicated in a broad range of cognitive functions, including memory consolidation, planning, and decision-making [10]. Importantly, replay extends beyond the passive reinstatement of past experiences. Replay has been observed during both sleep and wakefulness immediately following learning and has been hypothesized to support the flexible reorganization and recombination of previously encoded information, enabling the offline exploration of alternative behavioral trajectories and solution strategies [6, 11, 12]. These properties make replay particularly relevant to insight, which is widely believed to involve the restructuring of existing knowledge and the formation of novel associations. By promoting the formation of novel relationships among previously encoded memories, replay may provide a mechanistic link between incubation and the emergence of insight.

Directly testing this hypothesis, however, presents substantial methodological challenges, particularly in humans performing complex cognitive tasks. Replay events are inherently spontaneous, transient, and embedded within ongoing neural activity, making them difficult to identify without external markers [13]. Furthermore, replay often occurs in a temporally compressed form, creating a distributional mismatch between the stimulus-evoked neural responses used to train decoding models and the neural dynamics expressed during offline reactivation [11, 12]. Consequently, direct investigation of the role of replay in creative cognition and insight has remained limited. Recent advances in neural decoding have begun to address these challenges by enabling the detection of memory reactivation and replay from noninvasive electrophysiological recordings in humans using machine learning techniques [10, 12–14]. Despite these methodological advances, it remains unclear whether neural activity during incubation merely reflects memory replay or actively simulates problem related information in a manner that promotes insight and successful problem solving.

To investigate whether offline neural activity during incubation contributes to insight, a visual sequential insight task (VSIT), inspired by the number reduction task [15], was developed in which participants could discover a hidden rule that dramatically improved task performance. The paradigm was combined with EEG-based neural decoding to track the spontaneous reactivation of task representations during incubation and reconstruct their temporal organization. This approach enabled the examination of whether replay and simulation of problem-related information predict subsequent insight, as well as the characterization of the neural dynamics associated with successful problem solving.

## 2 Results

### 2.1 Visual sequential insight task (VSIT): a paradigm for decoding incubation

To investigate whether offline replay contributes to insight, we developed a visual sequential insight task (VSIT; Fig. 1A). In each trial, participants first viewed three response options, followed by the sequential presentation of two visual stimuli. They were instructed to predict the correct target as quickly and accurately as possible and could respond either after the first stimulus or, if additional information was needed, after the second stimulus. Immediate feedback was provided after each trial to facilitate learning. The VSIT was designed to elicit insight through the discovery of a hidden rule.

**Fig. 1:**
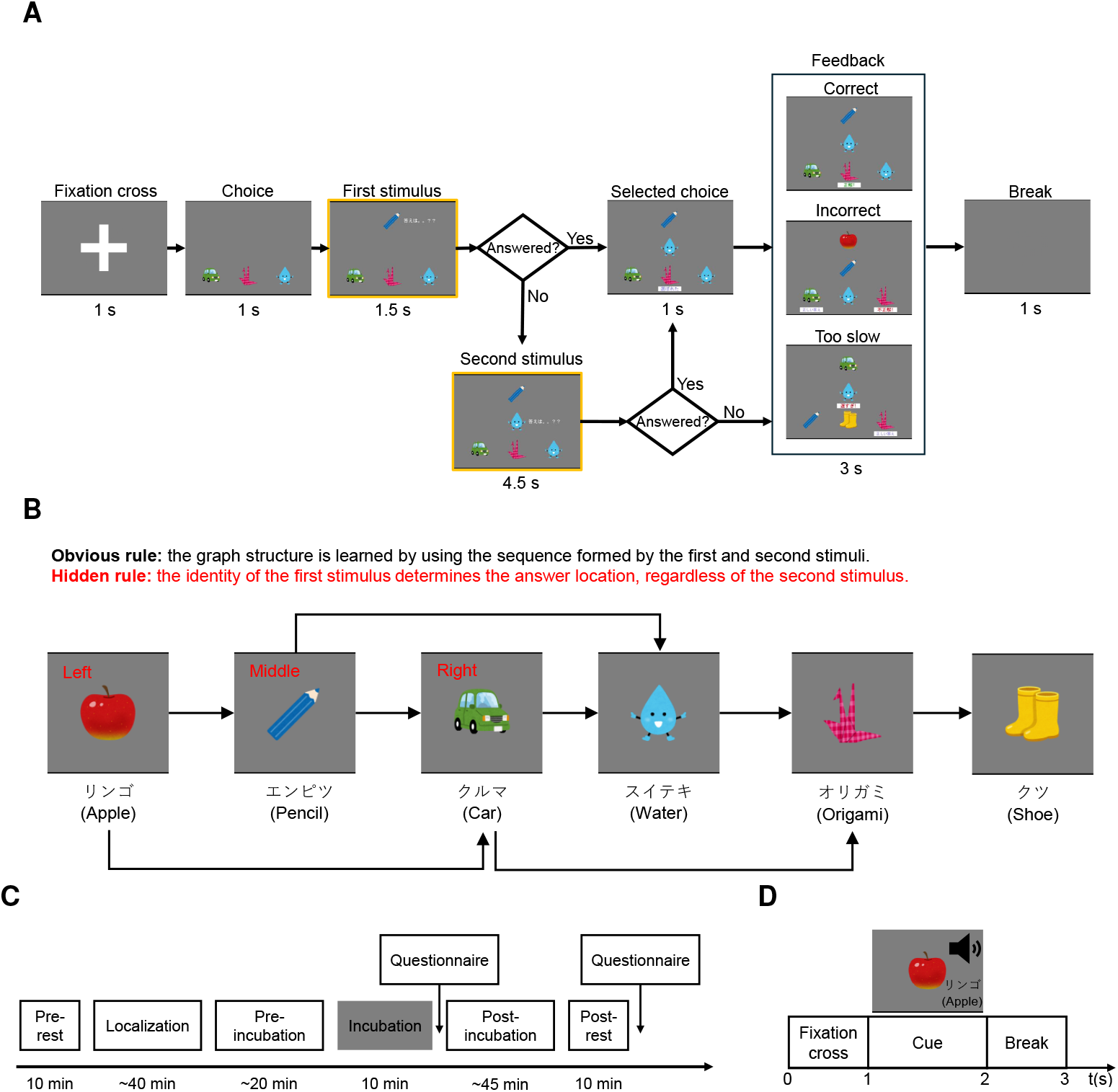
Task design and experimental protocol. **A** The visual sequential insight task (VSIT) trial structure, including the initial choice display, first stimulus onset, second stimulus onset, and the logic mechanism triggered by a response within the permitted window (yellow border). Feedback is provided at the end of every trial to facilitate associative learning. **B** Task rules underlying the VSIT include an obvious rule that requires the acquisition of sequential transitions (First → Second → Target) and a hidden rule that enables an immediate behavioral response based solely on the spatial mapping of the first stimulus. Specifically, the first stimulus directly indicates the location of the correct response target (e.g., “Apple” maps to the left, “Pencil” to the middle, and “Car” to the right). **C** Session structure, including resting-state EEG recordings (pre-, incubation, and post-rest), the localization session, and the two stages of VSIT conducted during the pre-incubation and post-incubation sessions. Questionnaires were administered directly after incubation and post-rest sessions to assess subjective strategy and insight. **D** Localization session, where participants performed a passive viewing and internal rehearsal task with auditory cues to establish baseline neural representations for the six target stimuli.

To facilitate subsequent decoding of memory reactivation, the task employed visually and auditorily distinct stimuli that maximized the discriminability of item-specific neural representations. The task contained an explicit rule that could be learned by memorizing six stimulus sequences and a hidden rule in which the first stimulus alone determined the correct response location (Fig. 1B). Once discovered, this shortcut allowed participants to respond before the second stimulus appeared, substantially improving performance.

The overall experimental timeline is shown in Fig. 1C. Participants completed the VSIT before (pre-incubation) and after (post-incubation) the incubation period. Excluding the training run, the pre-incubation session consisted of three runs (54 trials), whereas the post-incubation session consisted of twelve runs (216 trials). Each run comprised 18 trials, with each of the six sequences repeated three times. Insight was defined as successful discovery of the hidden rule and was determined using both objective behavioral criteria and post-experiment questionnaires (described in the next section).

To enable decoding of spontaneous memory reactivation during the incubation period, participants first completed a localization session to obtain stimulus-specific EEG responses associated with the task items (Fig. 1D). These responses were subsequently used to train neural decoders capable of identifying spontaneous memory reactivation events during the incubation period.

### 2.2 Behavioral trajectories reveal abrupt post-incubation transitions to insight

Representative behavioral trajectories from one participant in the no-insight group and one participant in the insight group are shown in Fig. 2A. Behavioral performance was quantified using a trial-wise composite score, defined as accuracy divided by reaction time. Because the second stimulus appeared 1.5 s after the first, a composite score exceeding 1/1.5 indicated that a participant had responded correctly before presentation of the second stimulus, demonstrating use of the hidden rule. Insight was therefore identified when the composite score transitioned from below to above this threshold, and remained above it for at least 60% of the subsequent 15 trials (see details in Methods). To further characterize individual learning trajectories, trial-wise composite scores were fitted with a Gompertz function (representative examples are shown in Fig. 2A, and fits for all participants are shown in Supplementary Fig. 1). From each fitted curve, two behavioral metrics were derived: the inflection point, corresponding to the maximum learning rate and interpreted as the onset of insight, and the maximum learning rate, corresponding to the maximum slope of the Gompertz function.

**Fig. 2:**
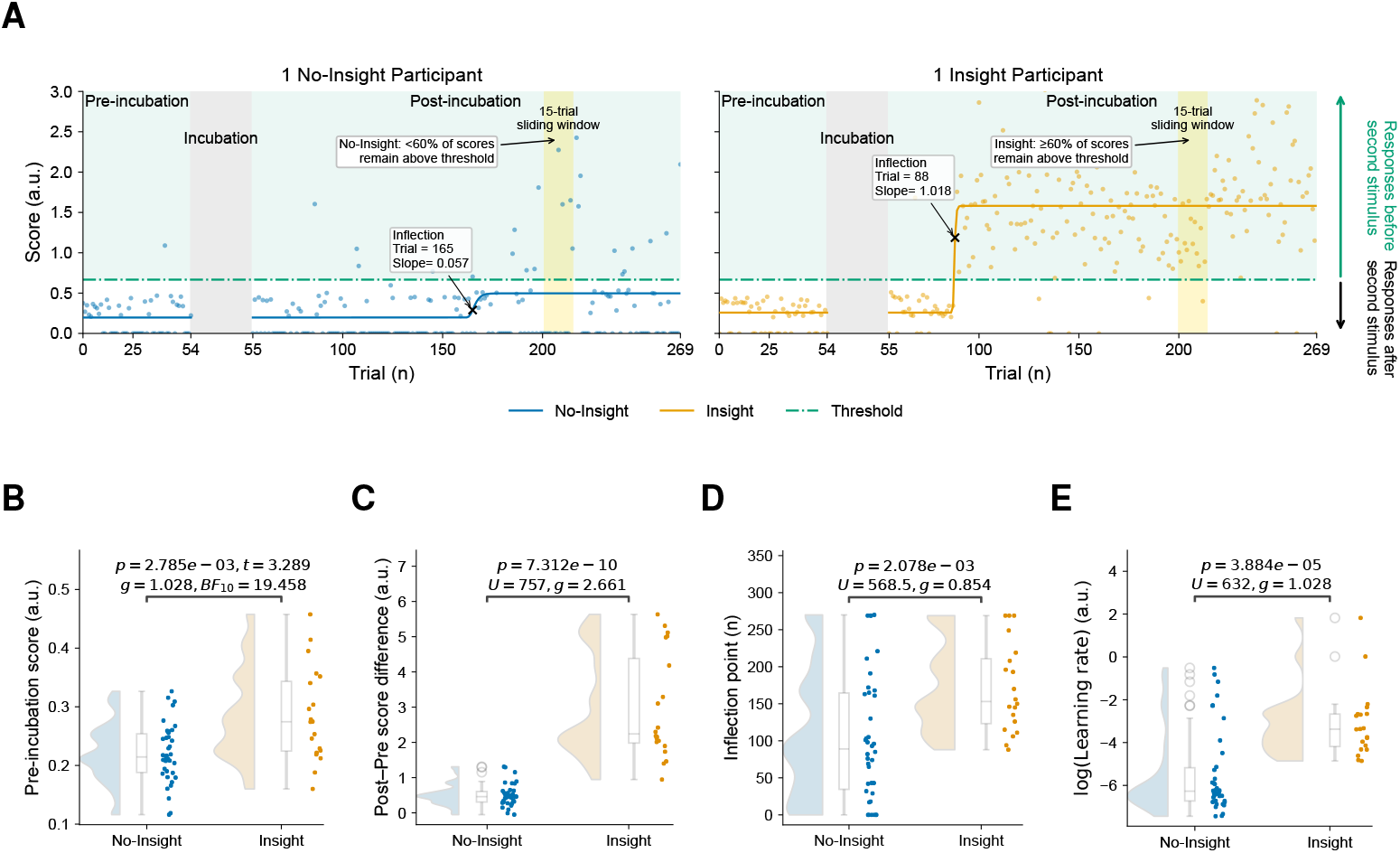
Behavioral differences between insight and no-insight groups. **A** Behavioral performance quantification. Representative score trajectories (circles) and corresponding Gompertz fits (solid lines) for one participant from the no-insight (blue) and insight (orange) groups. The gray shaded region indicates the incubation period. The horizontal green dash-dotted line denotes the predefined threshold (1*/*1.5), above which responses occurred before presentation of the second stimulus. Black crosses indicate the Gompertz inflection point, corresponding to the maximum learning rate or onset on insight. The yellow shaded region illustrates a representative 15-trial sliding window used for insight detection. Insight was identified when the score transitioned from below to above the threshold and remained above the threshold for at least 60% of the subsequent 15 trials. The insight group exhibited significantly higher values across all behavioral measures, including **B** pre-incubation score, **C** score change between sessions, **D** estimated inflection point corresponding to the onset of score saturation, and **E** learning rate. Exact statistical values for group comparisons—including the significance level (*p*), Mann–Whitney test statistic (*U* ), Student’s *t*-statistic (*t*), Hedges’ *g* effect size (*g*), and Bayes factor (BF_10_)—are displayed at the top of each respective panel.

Participants who satisfied this behavioral criterion and correctly described the hidden rule in the post-experiment questionnaire were classified as belonging to the insight group. Using these behavioral and questionnaire criteria, participants were classified into either the insight group (*n* = 20) or the no-insight group (*n* = 38). Although two participants in the no-insight group correctly described the hidden rule, they did not meet the full behavioral criteria of the insight classification algorithm (see Supplementary Fig. 2 for raw scores). The resulting groups were then compared on baseline measures, including sleep quality, intelligence quotient, daydreaming frequency, and mind-wandering tendency. No significant differences were observed (all *p >* 0.05), indicating comparable demographic and cognitive profiles (Supplementary Fig. 3).

The insight group exhibited significantly higher mean scores during the pre-incubation session (*p* = 2.785*e* −3; two-tailed Mann-Whitney *U* test; Fig. 2B), suggesting that participants who ultimately discovered the hidden rule learned the obvious rule more efficiently during initial exposure, potentially freeing cognitive resources for exploration of alternative strategies. Furthermore, participants in the insight group exhibited a significantly greater increase in score from the pre-incubation to the post-incubation session than those in the no-insight group (*p* = 7.312*e* −10; two-tailed Mann-Whitney *U* test; Fig. 2C), consistent with substantial performance gains following hidden-rule discovery.

The insight group also exhibited a significantly later Gompertz inflection point than the no-insight group (*p* = 2.078*e* − 3; two-tailed Mann-Whitney *U* test; Fig. 2D), indicating a more prolonged learning trajectory before reaching rapid improvement. This delayed inflection likely reflects the transition from learning the obvious rule to discovering the hidden rule, whereas the no-insight group reached performance saturation after acquiring only the obvious rule. In addition, the insight group demonstrated a significantly higher maximum learning rate (*p* = 3.884*e* − 5; two-tailed Mann-Whitney *U* test; Fig. 2E), indicating that discovery of the hidden rule produced a more abrupt improvement in performance.

### 2.3 Stable EEG decoding enables tracking of spontaneous memory reactivation

To examine what occurred during the incubation period that led to the greater performance improvement and faster learning observed in the insight group, spontaneous reactivation of task items (e.g., an apple) during incubation was decoded. A localizer model was trained on EEG data acquired during the localization session and subsequently applied to EEG recordings collected during incubation to decode spontaneous memory reactivation events. The decoded reactivation sequences were reconstructed into replay trajectories and analyzed using replay template-matching analyses (see the complete analysis pipeline in Fig. 3). Replay-derived features were then used to determine whether replay dynamics during incubation predicted subsequent insight.

**Fig. 3:**
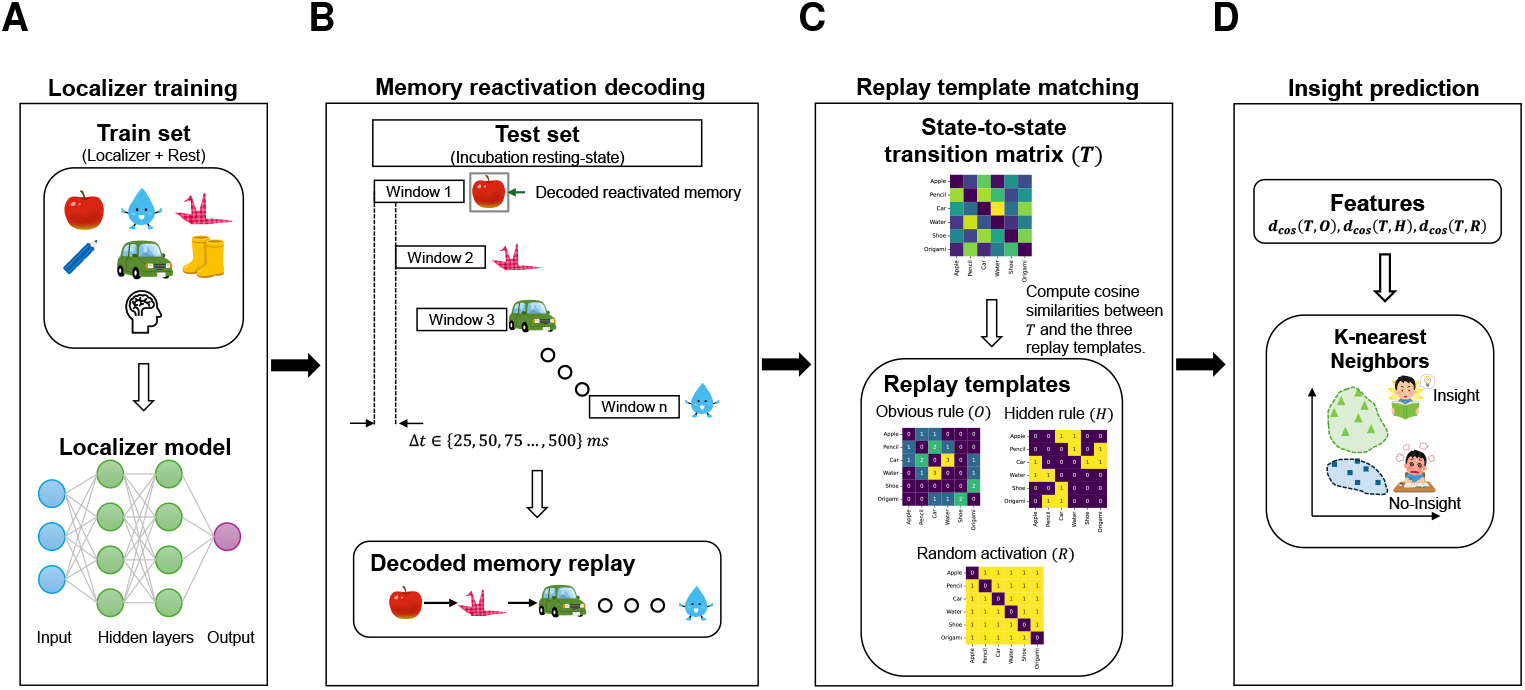
Analytical pipeline for insight decoding. **A** A localizer model was first trained to classify six image categories and resting-state activity from EEG recordings. **B** The optimized decoder was subsequently applied to incubation-session EEG data to identify spontaneous neural reactivation events. **C** Decoded reactivation sequences were then organized into empirical transition matrices and compared with predefined theoretical replay templates using cosine similarity measures to derive replay-structure features. **D** Finally, these replay features were used in binary classification to predict subsequent behavioral insight.

Accurate decoding of replay depends on the assumption that spontaneous memory reactivation resembles stimulus-evoked activity recorded during the localization session. However, replay events are often temporally compressed relative to the original experience, creating a distributional mismatch between the stimulus-evoked activity used for training and the neural activity observed during incubation [11, 16]. To mitigate this covariate shift, EEG-specific data augmentation techniques were incorporated during training (see Supplementary Table 1). Four state-of-the-art EEG decoding architectures, including EEGNet [17], ShallowConvNet [18], EEGITNet [19], and Conformer [20], were trained and evaluated using localization-session recordings comprising stimulus-evoked responses to VSIT images and resting-state data collected during the pre-rest session. Their decoding performance was compared to identify the optimal localizer model.

A linear mixed-effects model evaluating classification performance as a function of model architecture, behavioral group, and their interaction revealed a significant main effect of model architecture (*p* = 2.803*e* − 15). In contrast, the main effect of behavioral group (*p* = 0.783) and the interaction term (*p* = 0.475) were non-significant (detailed model parameters are provided in Supplementary Table 2). Subsequent post-hoc two-tailed Wilcoxon signed-rank tests with Bonferroni correction showed that EEGNet achieved the highest mean decoding accuracy (Fig. 4A). However, its performance did not differ significantly from that of ShallowConvNet.

**Fig. 4:**
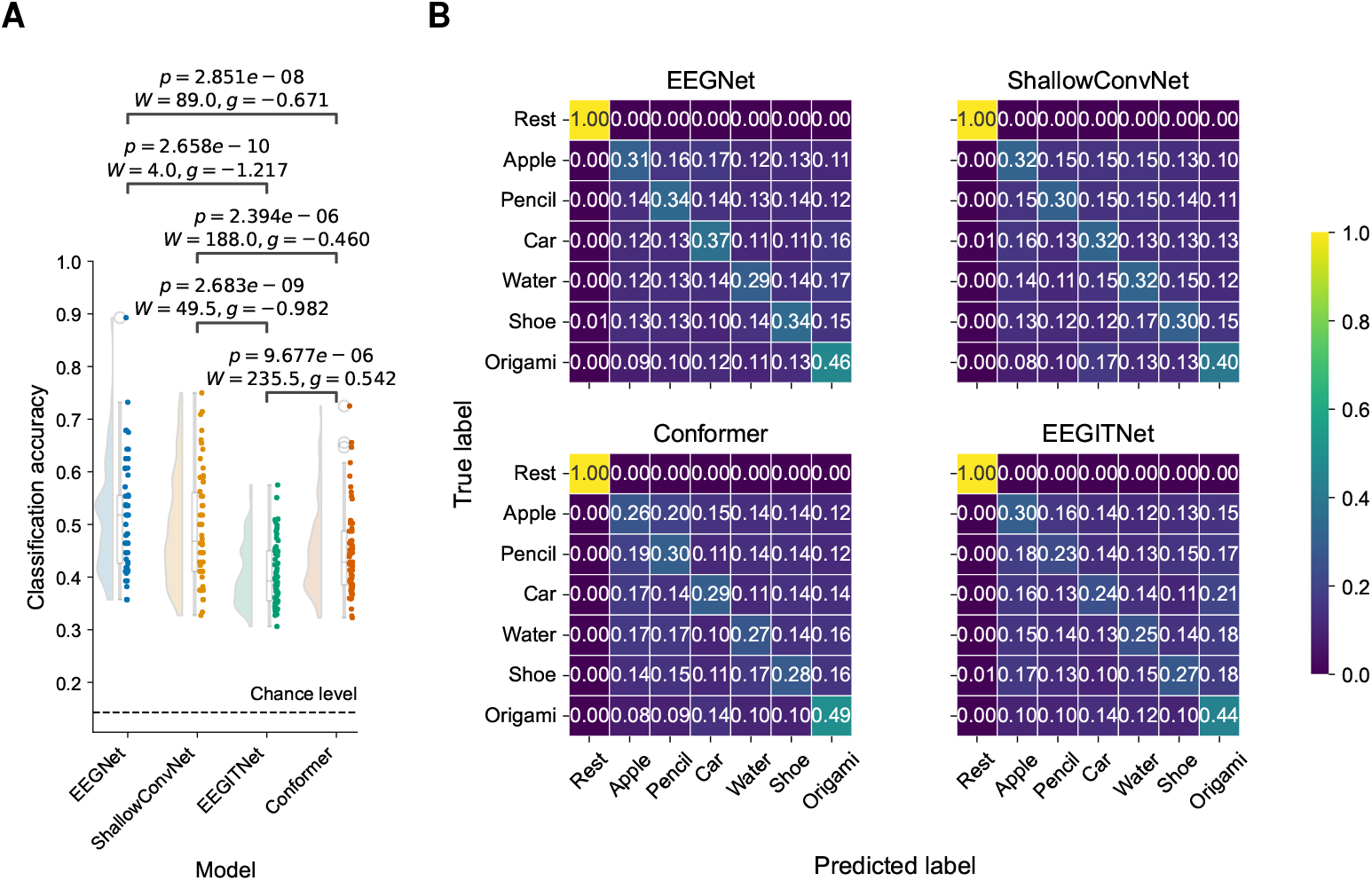
Comparison of localizer model performance. **A** Classification accuracy across localizer models. EEGNet achieved the highest decoding accuracy among all evaluated architectures (linear mixed-effects model; main effect of model: *p* = 2.803*e* − 15). Post-hoc statistical comparisons were performed using two-tailed Wilcoxon signed-rank tests with Bonferroni correction, with the corresponding *p*-values, Wilcoxon *W* statistics, and Hedges’ *g* effect sizes displayed directly above each model comparison. **B** Confusion matrices averaged across participants. Resting-state trials exhibited high classification accuracy, whereas task-related stimulus conditions showed moderate overlap between categories across all models.

Confusion matrices (Fig. 4B) further clarified model behavior, revealing that all classifiers exhibited similar performance patterns. Specifically, the models accurately identified the resting state, likely due to its distinct neural signature compared to task-related stimulus processing. In contrast, classification between different stimulus types showed higher confusion, suggesting possible shared neural features between localization stimuli. Therefore, EEGNet was selected as the localizer model for all subsequent replay decoding analyses because it achieved the highest overall classification performance. The generalization gap and calibration profiles of the models are detailed in the supplementary material (Supplementary Fig. 4 and 5).

### 2.4 Incubation-period memory replay predicts subsequent insight

The optimized EEGNet model was subsequently used to decode spontaneous memory reactivation during the incubation period. We next asked whether the structure of these replay events predicted which participants would subsequently experience insight. To quantify replay structure, decoded reactivation events were organized into replay sequences and compared with three predefined replay templates representing the obvious rule (*O*), the hidden rule (*H*), and random reactivation (*R*) (see details in Methods). These templates quantified the extent to which replay followed the learned task structure, the hidden solution strategy, or unstructured transitions.

Out-of-fold (OOF) prediction performance was evaluated across replay sequences constructed using multiple temporal lag conditions. Replay features derived using a 400 ms lag significantly predicted subsequent behavioral insight (*p*_uncorrected_ = 0.026; Fig. 5A). The corresponding precision-recall (PR) curve and non-parametric permutation test results are shown in Fig. 5B and Fig. 5C, respectively. The optimized decoding model achieved an area under the PR curve (PR-AUC) of 0.557, exceeding the chance baseline of 0.345 determined by the underlying class imbalance. This improvement indicates that replay structure during the incubation period contained information predictive of subsequent insight. Model prediction probabilities, calibration profiles, and receiver operating characteristic analyses are provided in Supplementary Fig. 6. Importantly, classifiers trained using only forward or only backward replay templates did not achieve above-chance performance (Supplementary Fig. 7), suggesting that successful prediction required information from both replay directions.

**Fig. 5:**
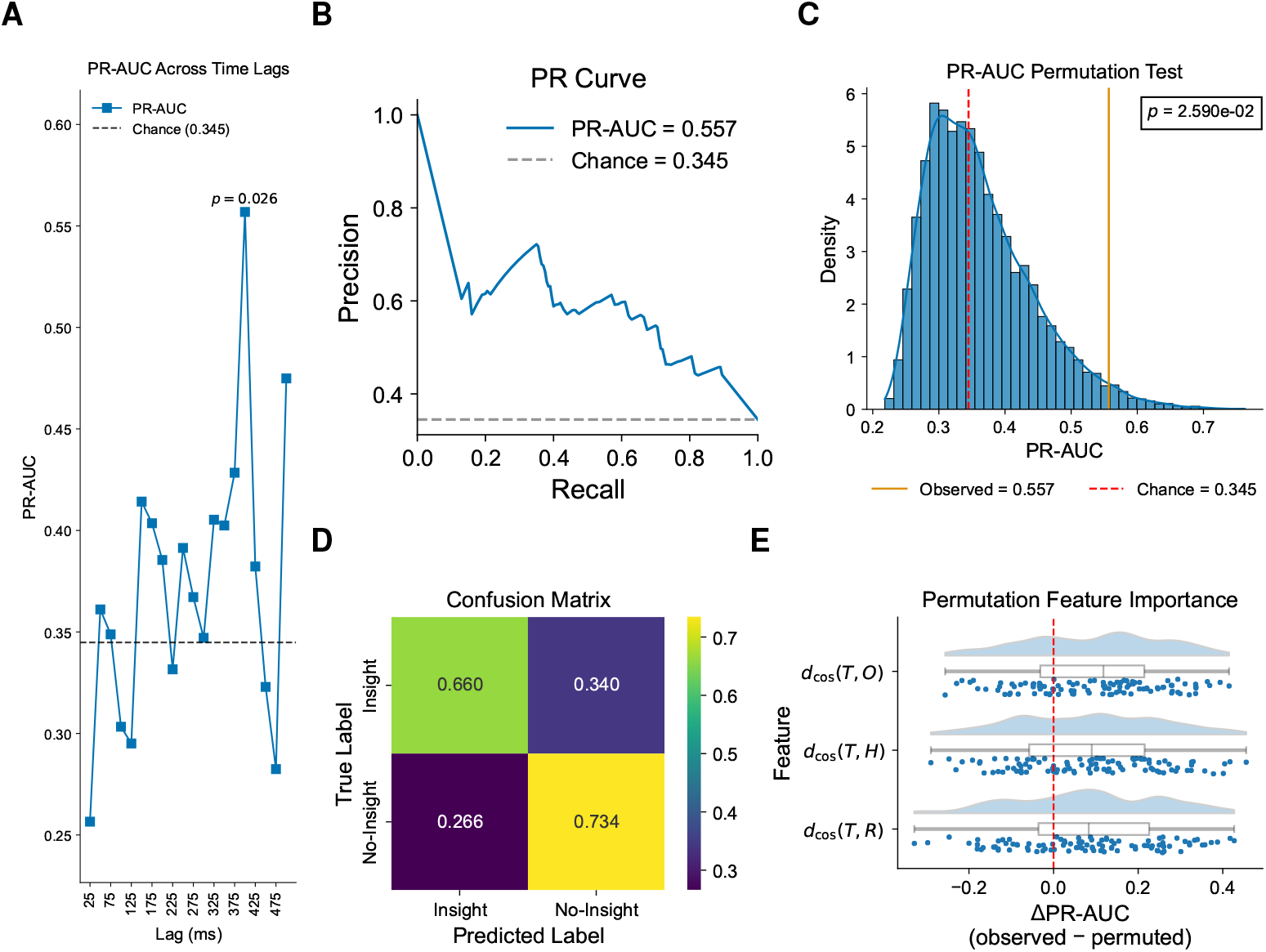
Insight prediction from 400 ms memory replay features using k-nearest neighbors model. **A** Classification performance across temporal lag conditions, quantified using area under the precision-recall curve (PR-AUC). Significant prediction performance was observed specifically for the 400 ms replay condition (permutation *p*_uncorrected_ = 0.026). **B** PR curve for the 400 ms condition, demonstrating reliable discrimination between insight and no-insight participants. **C** Null distribution obtained from 10,000 permutation tests for the 400 ms condition. **D** Normalized confusion matrix summarizing classification performance. **E** Permutation-based feature importance analysis, revealed a hierarchical contribution pattern: *d*_cos_(*T, O*) showed the greatest predictive importance, followed by *d*_cos_(*T, H*) and *d*_cos_(*T, R*).

The normalized OOF confusion matrix (Fig. 5D) further demonstrated reliable further demonstrated reliable classification performance, with a recall of 0.660, a specificity of 0.734, and an *F*_1_-score of 0.610. Permutation-based feature importance analysis revealed that all replay templates contributed to insight prediction (Fig. 5E). Among these, similarity to the obvious-rule template (*d*_cos_(*T, O*)) contributed most strongly to classifier performance, followed by the hidden-rule (*d*_cos_(*T, H*)) and random-replay (*d*_cos_(*T, R*)) templates.

Collectively, these findings demonstrate that structured memory replay during the incubation period predicts subsequent insight. Moreover, successful prediction required the integration of both forward and backward replay dynamics, suggesting that bidirectional replay contains complementary information relevant to the emergence of insight

### 2.5 Insight probability depends on higher-order interactions between replay structures

Although the preceding analysis demonstrated that replay structure predicted subsequent insight, it did not reveal how different replay structures contributed to this prediction. Because the k-nearest neighbors classifier does not provide directly interpretable feature weights, a post-hoc ordinary least squares (OLS) regression was performed to characterize the relationship between replay features and the classifier’s predicted probability of insight (Fig. 6; see Methods for details). Replay similarity to the obvious-rule, hidden-rule, and random-replay templates, together with their interaction terms, were included as predictors. The fitted model explained a substantial proportion of the variance in predicted insight probability (*R*^2^ = 0.412, *F*_7,50_ = 9.478, *p* = 1.989*e* − 7), indicating that the replay features captured meaningful variation in the classifier output.

**Fig. 6:**
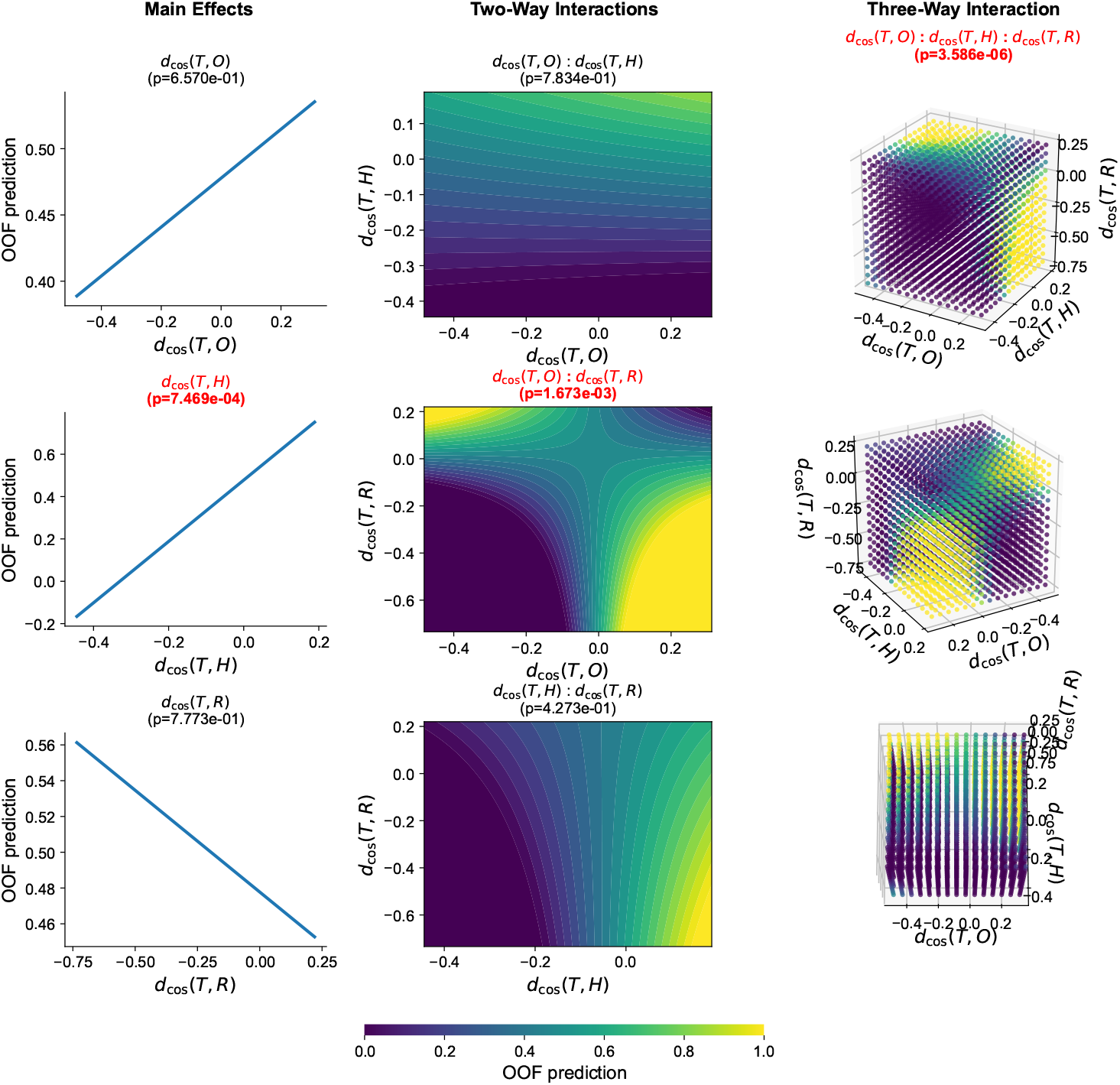
Modeling of insight prediction from memory replay structures. Ordinary least squares (OLS) regression analysis evaluated the relationship between mean out-of-fold (OOF) prediction probabilities and structural replay-similarity features, including their interactions. Statistically significant effects (*p <* 0.05) are highlighted in red. (Left) Main effect analysis reveals a significant isolated contribution of *d*_cos_(*T, H*). (Middle) Two-way interaction contour plots illustrate how pairs of replay-similarity features jointly modulate the predicted probability of subsequent insight. A significant interaction effect is observed for *d*_cos_(*T, O*) : *d*_cos_(*T, R*). (Right) Three-way interaction slice visualizations depict the complex combined influence of *d*_cos_(*T, O*), *d*_cos_(*T, H*), and *d*_cos_(*T, R*) across the complete multi-dimensional feature space.

Analysis of the main effects revealed that only replay similarity to the hidden-rule template (*d*_cos_(*T, H*)) significantly increased the predicted probability of insight (*p* = 7.469*e* − 4) (Fig. 6, left), suggesting that replay consistent with the hidden rule is a key contributor to subsequent insight.

Two-way interaction analyses further revealed a significant interaction between replay similarity to the obvious-rule and random-replay templates (*d*_cos_(*T, O*) : *d*_cos_(*T, R*); *p* = 1.673*e* − 3) (Fig. 6, middle). High predicted insight probability occurred under two complementary replay configurations: high similarity to the random template together with low similarity to the obvious-rule template, or conversely, high similarity to the obvious-rule template together with low similarity to the random template.

Finally, a significant three-way interaction among replay similarity to the obvious-rule, hidden-rule, and random-replay templates (*d*_cos_(*T, O*) : *d*_cos_(*T, H*) : *d*_cos_(*T, R*); *p* = 3.586*e* −6) further refined this relationship (Fig. 6, right). The model predicted the highest probability of insight under two distinct replay configurations: (1) high similarity to both the hidden-rule and random-replay templates, together with low similarity to the obvious-rule template; or (2) high similarity to both the hidden-rule and obvious-rule templates, together with low similarity to the random-replay template.

Collectively, these findings suggest that replay associated with the hidden rule is necessary but not sufficient to predict subsequent insight. Instead, insight probability depends on higher-order interactions among replay structures, suggesting that successful incubation reflects a dynamic balance between replay of the learned task structure and more exploratory replay patterns.

### 2.6 Insight is associated with enhanced low-gamma network integration

Having established that replay dynamics during incubation predicted subsequent insight, we next asked whether these replay-related processes were accompanied by differences in large-scale functional brain networks. Source-level functional connectivity analysis revealed a significant subnetwork exhibiting stronger low-gamma (30—50 Hz) functional connectivity in the insight group than in the no-insight group (*p* = 0.042) (Fig. 7). Functional connectivity was quantified using the weighted phase-lag index (wPLI) [21] and statistically evaluated using the network-based statistic (NBS) (see Methods for details).

**Fig. 7:**
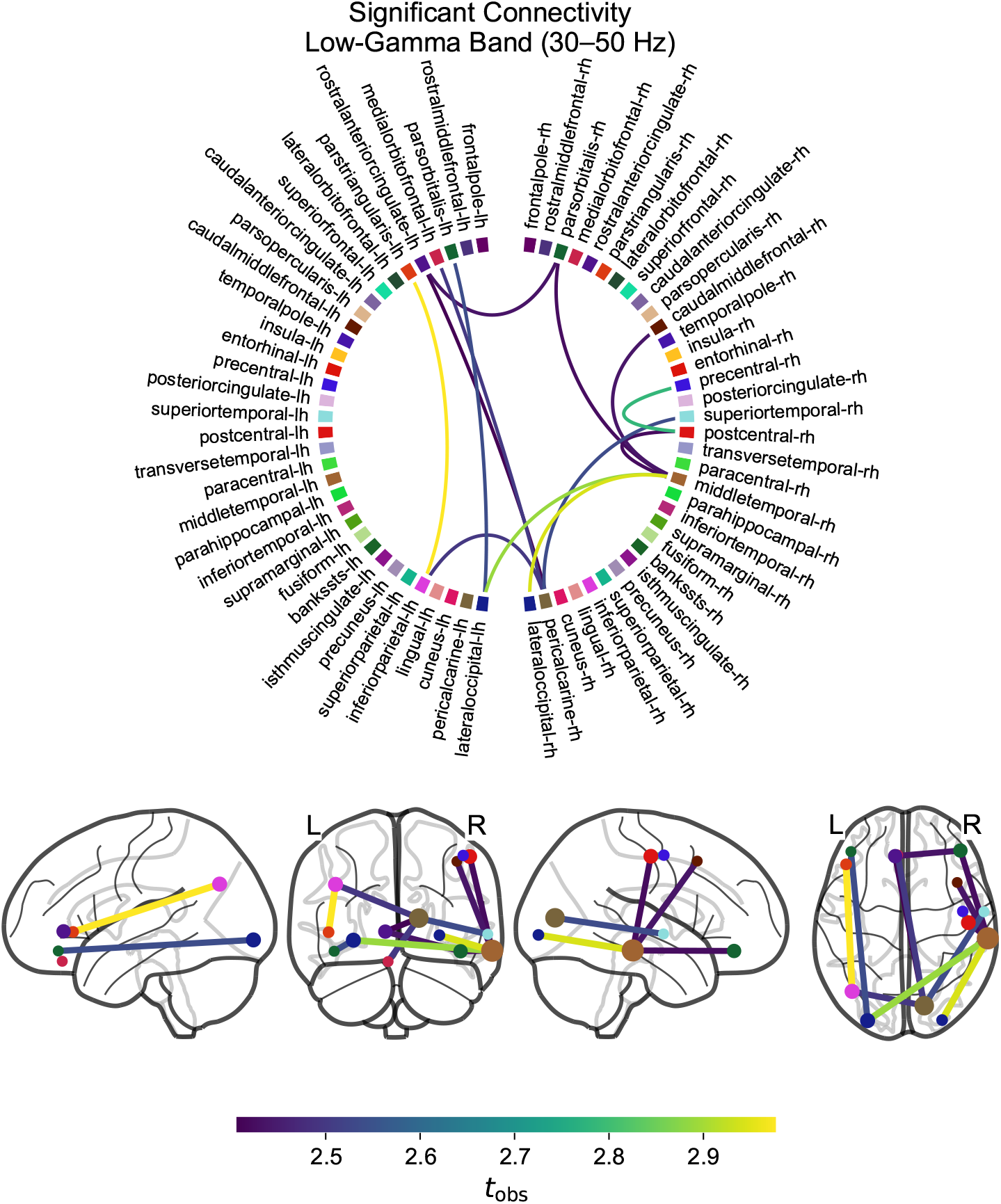
Source-level functional connectivity differences between groups in the low-gamma band (30–50 Hz). The top panel shows a connectome circle plot illustrating the significant subnetwork exhibiting stronger functional connectivity in the insight group relative to the no-insight group (*p* = 0.042). The bottom panel presents the corresponding anatomical projections on the cortical surface. For visualization purposes, only nodes with significant connections are displayed, and node size is proportional to degree (i.e., the number of significant connections). Edge colors represent the observed *t*-statistics (*t*_obs_).

The identified subnetwork was centered on two hub regions: the right middle temporal gyrus, a region implicated in memory retrieval and semantic integration, and the right pericalcarine cortex, which is primarily involved in visual processing. These hubs were connected to a distributed set of regions associated with semantic processing (pars orbitalis, pars triangularis, and superior temporal cortex), reward anticipation and value representation (orbitofrontal cortex), working memory and executive control (caudal middle frontal cortex), memory consolidation and retrieval (temporal regions), visual processing (pericalcarine and occipital cortices), and sensorimotor processing (precentral and postcentral gyrus).

This widespread increase in low-gamma synchronization suggests that the integration of task-relevant information during incubation is supported by coordinated communication across functionally specialized cortical systems. Such a distributed network architecture may reflect goal-directed mental simulation of task-related representations, facilitating the memory restructuring and associative recombination processes necessary for successful solution of the VSIT.

## 3 Discussion

The present study provides evidence that spontaneous memory replay during incubation contributes to the emergence of insight. By decoding replay from resting-state EEG, we found that replay dynamics predicted subsequent discovery of a hidden rule and were accompanied by enhanced low-gamma functional connectivity across distributed cortical networks. These findings suggest that incubation is not a passive period, but an active process in which replay reorganizes task representations to facilitate creative problem-solving.

Previous research has often relied on subjective “Aha!” ratings as the primary measure of insight. However, these ratings are susceptible to reporting bias and may reflect false insight rather than correct solution discovery [22]. By contrast, the present study combined objective behavioral performance with post-experiment questionnaires to classify participants into insight and no-insight groups, providing a more robust assessment of insight.

A central theoretical question of this study is how subconscious spontaneous memory replay during incubation contributes to insight. Although it is well established that taking a break from a problem or engaging in an unrelated task facilitates insight, the neurocognitive mechanisms by which subconscious processing promotes subsequent conscious problem-solving remain poorly understood [3, 4]. One possible mechanism is that subconscious replay actively reconstructs and reorganizes stored representations in a goal-directed manner, enabling previously unrecognized latent regularities to emerge [10]. Converging electrophysiological evidence suggests that awake human memory replay occurs in both temporal directions relative to the originally encoded experience, with each direction supporting distinct cognitive functions [12, 23, 24]. Specifically, forward replay has been linked to memory consolidation and the stabilization of learned representations, whereas backward replay has been associated with reward processing [16, 25, 26]. Consistent with this account, subsequent insight was predicted only when replay templates incorporated both forward and backward transitions. This finding suggests that effective incubation depends on the interaction of bidirectional replay processes. Forward replay may reinforce the obvious-rule representation, whereas backward replay may promote exploration of alternative pathways and latent solution strategies, facilitating access to the hidden rule.

In addition to replay direction, it is also important to consider the structure of replay events that facilitate insight. The interaction between the default mode network (DMN) and the executive control network (ECN) provides one possible neurocognitive framework for understanding these replay dynamics [27, 28]. According to this framework, the DMN supports the spontaneous recombination of stored memory traces to generate candidate representations, whereas the ECN exerts top-down control processes that evaluate, maintain, and prioritize task-relevant representations. Consistent with this framework, modeling insight probability from memory replay features revealed higher-order interactions among replay templates. In particular, greater replay similarity to the hidden-rule template (*d*_cos_(*T, H*)) significantly predicted subsequent insight, suggesting that replay may represent latent task structure before it becomes consciously accessible. Through coordinated interactions between the DMN and ECN, replayed memory representations may be progressively refined until latent regularities become integrated into an explicit solution, ultimately giving rise to insight.

Additionally, higher insight probability was associated with two distinct replay configurations: (1) high similarity to both the hidden-rule and obvious-rule (*d*_cos_(*T, O*)) templates together with low similarity to the random template (*d*_cos_(*T, R*)), and (2) high similarity to the hidden-rule and random templates together with low similarity to the obvious-rule template. Viewed through the DMN–ECN framework, the first configuration may reflect a state in which executive control maintains replay within task-relevant representational boundaries while allowing explicit and latent task structures to be concurrently reactivated. Such coordinated replay may facilitate the integration and restructuring of existing knowledge, enabling previously unrecognized relationships between the obvious and hidden rules to emerge [29]. In contrast, the second configuration may reflect a more exploratory state characterized by reduced dominance of the obvious-rule representation and greater engagement of less constrained replay dynamics. This interpretation is consistent with evidence that creativity emerges through exploration of less-traveled representational trajectories [8, 30–32] and that increased neural stochasticity enhances creative performance [33]. Although no group differences in self-reported mind wandering were observed, replay resembling the random template may nevertheless promote exploration of alternative associations and the recombination of previously unrelated information into novel solutions. Notably, both configurations were characterized by strong hidden-rule replay, suggesting that reactivation of latent task structure is a common prerequisite for insight. The accompanying replay dynamics, however, may determine whether insight emerges through structured integration of explicit and latent task representations or through exploratory processing that relaxes constraints imposed by dominant solution strategies [15]. Together, these findings suggest that successful incubation depends on balancing exploitation of established knowledge with exploration of alternative representational configurations [34]. Such a balance may arise through coordinated interactions between the DMN and ECN, enabling memory replay to both stabilize existing knowledge and generate novel representational pathways that culminate in insight [27, 28].

Furthermore, source-space network analyses revealed significant group differences in low-gamma functional connectivity involving regions associated with visual processing, memory, reward, and sensorimotor functions, consistent with previous studies linking creativity to distributed neural networks [35, 36]. Gamma-band activity has been implicated in integrating distributed information, coordinating sensory and memory processes, and supporting conscious awareness [37–39]. Enhanced low-gamma connectivity during incubation may therefore reflect the subconscious integration and reorganization of memory representations that support subsequent insight. This interpretation is consistent with evidence linking gamma oscillations to awake hippocampal replay, where transient synchronization coordinates memory reactivation and information transfer [10, 40]. Previous studies have also associated insight with transient gamma-band activity in the right temporal cortex and enhanced temporal connectivity during creative cognition [9, 29, 41–43]. Consistent with these findings, a temporal cortical region emerged as a central hub in the present low-gamma network. Notably, whereas previous studies have focused on neural activity immediately preceding or following insight [38], the connectivity differences observed here occurred during incubation, before conscious solution awareness. These findings suggest that gamma-mediated processes supporting insight begin during incubation, reflecting the progressive integration and replay-driven reorganization of memory representations prior to conscious solution emergence.

Despite the present findings demonstrating that spontaneous memory replay contributes to insight, replay alone is unlikely to fully account for insight. Although insight is generally thought to emerge through the restructuring of problem representations, attentional, emotional, and executive control processes are also likely to play important roles [2, 7]. For example, solving the classic nine-dot problem requires shifting attention beyond an implicitly imposed boundary, whereas recognizing a meaningful image in an ambiguous arrangement of blobs requires perceptual restructuring. These examples suggest that memory replay alone is insufficient to explain insight, as successful problem solving also requires overcoming attentional fixation and relaxing constraints imposed by existing problem representations. In particular, mental set and functional fixedness bias attention toward familiar solutions, limiting the exploration of alternative representations [44]. Attentional flexibility, emotional processes, and executive control may help overcome these constraints by promoting exploration of alternative solution pathways and enabling reorganized representations to become consciously accessible as insight. Furthermore, the relative contribution of these processes may vary across insight domains. Whereas perceptual insight may rely more heavily on attentional and perceptual restructuring, semantic or knowledge-based insight may depend more on memory reactivation and associative recombination. Future studies comparing replay dynamics across diverse insight paradigms will help determine the extent to which the present findings generalize across different forms of insight.

A key methodological limitation of the present study is the lack of a ground-truth measure against which decoded memory reactivation can be validated. As all subsequent analyses were derived from these decoded replay events, the extent to which the identified patterns reflect true neural replay remains uncertain. Moreover, modeling insight probability from replay features provides only an indirect approximation of the neural processes underlying insight emergence. Although the observed associations between replay features and insight probability suggest potential mechanisms linking replay and creative problem-solving, they remain correlational and do not establish causality. Importantly, the predictive mapping between replay features and insight was also examined using a surrogate modeling approach, which approximates the behavior of the primary decoding model to aid interpretability; however, its inferences reflect an approximation of the learned decision function rather than direct statistical effects in the neural data. Finally, although EEG offers excellent temporal resolution for tracking replay dynamics, its limited spatial resolution constrains precise source localization, and therefore source-space connectivity findings should be interpreted with caution. Multimodal approaches such as simultaneous EEG–fMRI may help address this limitation.

## 4 Methods

### 4.1 Participants

Sixty-four native or highly proficient Japanese speakers were recruited for the study (37 males, 27 females; mean age 24.95 ± 5.70 years). Inclusion criteria were an age range of 18–40 years, no history of neurological or psychiatric disorders, no current use of psychotropic medication, and no severe visual, motor, or auditory impairments. Individuals who were pregnant or reported claustrophobia were excluded. Data from six participants were excluded from the final analysis: two due to prior participation in the pilot study, three for discovering the hidden rule during the pre-incubation session, and one for failing to adhere to task instructions.

The insight group (*n* = 20) had a mean age of 23.75 ± 4.09 years and comprised 12 males and 8 females (18 right-handed, 2 left-handed), whereas the no-insight group (*n* = 38) had a mean age of 25.58±6.35 years and comprised 25 males and 13 females (36 right-handed, 1 left-handed, and 1 ambidextrous).

All participants provided written informed consent prior to the experiment. The protocol was approved by the Ethics Committee of the University of Tokyo (IRB Number: E2025ALS288) and conducted in accordance with the Declaration of Helsinki. Participants were informed that the experiment duration would be approximately three hours and were compensated at a rate of 1,000 JPY per 30 min.

### 4.2 Experimental protocol

Participants were seated in a soundproof chamber. The experimental paradigm was developed and executed using `PsychoPy` (v2025.2.3) [45], comprising six distinct sessions structured chronologically as illustrated in Fig. 1C.

#### 4.2.1 Resting-state recordings

Three 10-minute resting-state recordings were acquired: pre-rest, incubation and post-rest sessions. During these sessions, a fixation cross was presented on the screen. Participants were instructed to keep their eyes open, remain relaxed, and avoid engaging in directed mental activities or systematic thought.

#### 4.2.2 Localizer task

A localizer task was conducted to obtain EEG signatures for stimuli used in the VSIT (Fig. 1D). Each trial began with a 1 s fixation cross, followed by a 1.5 s bimodal stimulus presentation pairing a visual image with a simultaneous Japanese auditory word cue. Six distinct stimulus conditions varying in color, shape, and semantic category were employed; their Japanese labels utilized distinct initial phonetic sounds to maximize separability of their EEG representations (Fig. 1B). Each trial concluded with a brief rest interval. Participants completed eight runs, with each run containing 60 trials (10 trials per condition) presented in a randomized sequence. These data served as ground-truth labels for training the decoding models used for subsequent memory reactivation decoding during the incubation session.

#### 4.2.3 Visual sequential insight task (VSIT)

The VSIT was developed to investigate the transition from trial-and-error associative learning to rule-based insight. The temporal structure of the VSIT is illustrated in Fig. 1A. The task required participants to identify a target stimulus based on a sequence of preceding stimuli. Participants were instructed only to respond as quickly and accurately as possible.

The task was governed by two embedded rules (Fig. 1B):

1. **Obvious Rule:** Stimulus sequences followed an underlying deterministic graph structure consisting of six unique transition patterns. The first stimulus always belonged to one of three categories (e.g., “Apple,” “Pencil,” or “Car”). Because stimuli were associated with multiple potential transitions, participants typically had to wait for the second stimulus to reliably determine the correct answer. Learning this rule therefore relied on exploiting the sequential transition structure through repeated exposure across trials. To ensure that each first-stimulus category appeared equally often, the transition sequence Apple → Car → Origami was removed, resulting in each first stimulus occurring exactly twice.
2. **Hidden Rule:** The identity of the first stimulus directly indicated the spatial location of the correct response, with each category corresponding to a fixed response position (e.g., “Apple” mapped to the left, “Pencil” to the middle, and “Car” to the right). Successful discovery of this rule required exploratory search for an alternative solution rather than continued reliance on the obvious rule. Once identified, it allowed participants to respond immediately after the first stimulus, bypassing the second stimulus entirely.

Each trial began with a 1 s fixation cross, followed by the presentation of three response options at the bottom of the screen for 1 s. To prevent participants from solving the task via simple elimination, these options always included two predetermined target stimuli based on the graph structure and one random distractor. For example, if the “Apple” stimulus served as the first stimulus, it could lead to the potential targets “Car” or “Water” depending on the subsequent second stimulus (e.g., Apple → Pencil → **Car**; Apple → Car → **Water**). In this scenario, “Car” and “Water” were presented as options alongside a third randomly selected distractor.

The first stimulus then appeared at the top of the screen for 1.5 s; participants were permitted to register a response at any point following this onset. Subsequently, the second stimulus appeared in the middle of the screen for 1.5 s. If no response was registered by this point, an additional 3 s response window followed. In Fig. 1A, the period during which participants were permitted to register a response is indicated by an orange border. Upon registration of an answer, the sequence immediately advanced to a “Selected Choice” confirmation screen. Each trial included a 3 s feed-back period (e.g., “Correct,” “Incorrect,” or “Too Slow”). In cases of an incorrect or missing response, the correct target stimulus was displayed during this feedback period to facilitate learning. Each trial ended with a 1 s break.

Participants performed the VSIT across two sessions: a pre-incubation session (4 runs) and a post-incubation session (12 runs). Each run consisted of 18 trials, with each of the six patterns appearing three times per run. To ensure that participants understood the asynchronous nature of the task, the initial run of the pre-incubation session served as a training phase and was not used for analysis. During this phase, participants were specifically instructed to attempt answering approximately half of the trials immediately after the first stimulus appeared and the remaining half after the onset of the second stimulus.

Responses were registered via a standard keyboard, using the `F` and `J` keys for the left and right options, respectively, and the `Space` bar for the middle option. To maintain high levels of engagement, participants were informed that the experiment would terminate early upon successful task completion while still providing full monetary compensation for the full three-hour duration.

#### 4.2.4 Questionnaires

Prior to the experiment, participants completed a set of questionnaires, including the Flinders Handedness Survey [46], the Pittsburgh Sleep Quality Index [47], the Day-dreaming Frequency Scale [48], the Mind-Wandering Questionnaire [48], and Raven’s Progressive Matrices [49]. Task engagement, worry, and distress were subsequently assessed using the Short Stress State Questionnaire (SSSQ) [50] following both the incubation and post-rest sessions. Conscious rule discovery was evaluated via a questionnaire requiring participants to report any perceived patterns and provide detailed descriptions of identified rules.

### 4.3 Behavioral analysis

Behavioral performance was quantified using a trial-wise composite efficiency score:

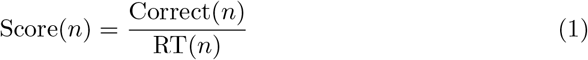

where Correct(*n*) ∈ {0, 1} and RT(*n*) represents the reaction time in seconds. RTs were calculated relative to the onset of the first stimulus. Because the second stimulus appeared 1.5 s after the first, a score exceeding 1*/*1.5 (0.667) indicated a correct response prior to the onset of the second stimulus.

Behavioral insight was identified by combining algorithmic detection and self-report. Algorithmic insight was defined as an abrupt and sustained increase in performance scores. Specifically, an insight event was logged at trial *n* if the score transitioned from below to above the threshold and remained above this threshold for at least 60% of the subsequent 15 trials. This objective criterion was validated against a post-experiment questionnaire in which participants reported the rule they believed they had identified. Participants were classified into the insight group only when algorithmic detection was consistent with participants’ self-reported discovery of the hidden rule.

Individual learning dynamics were fitted using a Gompertz growth function:

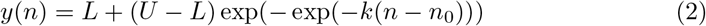

where *y*(*n*) denotes the predicted score at trial *n, L* and *U* represent the lower and upper asymptotes, and *k* determines the growth rate. The inflection point, *n*_0_, was utilized as a mathematical proxy for the onset of insight, representing the trial of maximal performance acceleration. The peak rate of change of performance improvement was quantified as the maximum learning rate (LR_max_):

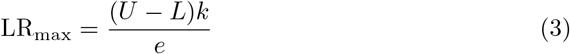

### 4.4 EEG acquisition and preprocessing

EEG signals were recorded using an active Ag/AgCl 64-channel electrode system. The signals were amplified utilizing a BrainAmp amplifier system (Brain Products GmbH, Gilching, Germany) at a sampling rate of 1 kHz. The ground and reference electrodes were positioned at channels FPz and FCz, respectively. Electrode impedance was periodically monitored throughout the recording sessions and maintained below 25 kΩ. To minimize physiological signal contamination and artifact, participants were instructed to maintain fixation and limit excessive eye blinking, head movements, and jaw clenching during the experiment.

The EEG data were band-pass filtered between 1 and 100 Hz and notch-filtered at 50 Hz to remove power-line noise. Filtering was implemented using zero-phase finite impulse response (FIR) filters with a Hamming window (0.0194 passband ripple, 53 dB stopband attenuation), utilizing a one-pass, non-causal design. Bad channels were identified through a combination of visual inspection and the local outlier factor algorithm. Independent component analysis was subsequently performed to identify and remove ocular and muscular artifacts. Following artifact rejection, any previously identified bad channels were reconstructed using spherical spline interpolation. All preprocessing steps were implemented using `MNE-Python` (v1.11.0) [51, 52].

### 4.5 Data analysis

An overview of the analysis pipeline is depicted in Fig. 3.

#### 4.5.1 Localizer training

This study evaluated four established EEG classification architectures—EEGNet [17], ShallowConvNet [18], EEGITNet [19], and Conformer [20]—as localizer models to decode memory reactivation during the incubation period. Detailed specifications regarding model architectures and hyperparameter configurations are provided in Supplementary Tables 4, 5, 6, and 7. The models were trained on data from localization sessions encompassing seven distinct classes: six image-cue conditions and one resting-state condition.

Additional signal processing involved high-pass filtering the raw EEG data at 4 Hz using a one-pass, zero-phase, non-causal FIR filter (Hamming window, 0.0194 pass-band ripple, 53 dB stopband attenuation, transition bandwidth: 2.00 Hz). The data were subsequently resampled to 250 Hz.

For each participant, EEG epochs were extracted within a temporal window of [−0.5, 1.0] s relative to stimulus onset. Artifact rejection was performed by excluding any epochs with peak-to-peak voltage amplitudes exceeding 150 *µ*V. To address potential class imbalance, event counts were equalized across conditions via random undersampling prior to model training. Following epoch extraction, exponential moving standardization was applied to the data.

Model training was performed using the Adam optimizer (learning rate = 5*e* −4, batch size = 32) for a maximum of 100 epochs. Prior to augmentation, EEG epochs were segmented into overlapping 1 s sliding windows with a 100 ms stride. To improve robustness against inter-individual EEG variability and unknown memory-reactivation distributions, extensive data augmentation was applied while preventing data leakage between training and validation sets (Supplementary Table 1). Overfitting was mitigated using early stopping with a patience of 20 epochs. Model performance was evaluated using stratified 10-fold cross-validation, and the model with the lowest validation loss for each participant was retained for subsequent analyses. All analyses were implemented in `PyTorch` (v2.10.0+cu130) [53] using the `Braindecode` library (v1.3.2) [54].

#### 4.5.2 Memory reactivation decoding

Following localizer training, the optimized model was applied to decode memory reactivation during the incubation session. The incubation EEG data underwent a preprocessing pipeline identical to that of the localization data.

EEG recordings obtained during the incubation session were segmented into overlapping 1 s sliding windows. Memory reactivation decoding was initially performed with a temporal stride of 25 ms to generate replay sequences. Additional lag conditions (25–500 ms, in 25 ms increments) were subsequently generated via temporal subsampling to examine the effect of temporal replay resolution on insight predictions.

#### 4.5.3 Empirical transition matrix construction and template matching

Decoded memory reactivations were mapped onto a state-transition framework to characterize replay dynamics during the incubation period. Let *S*_raw_ = (*s*_1_, *s*_2_, …, *s*_*N*_ ) represent the chronological sequence of maximum-probability state classifications across sliding windows, where *s*_*t*_ ∈ {0, 1, …, 6} corresponds to either a resting-state baseline (0) or one of six task-related image categories (1 ≤ *s*_*t*_ ≤ 6: Apple, Pencil, Car, Water, Shoe, or Origami). Memory reactivations were isolated by excluding all baseline resting-state predictions (*s*_*t*_ ≠ 0), and the remaining category indices were shifted down to define an operational state-space *X* = {0, 1, …, 5 }.

Transitions between distinct representational states were emphasized by compressing consecutive repetitions of identical states into a state-change sequence *S*_collapsed_ = (*c*_1_, *c*_2_, …, *c*_*M*_ ), where *c*_*m*_ ≠ *c*_*m*+1_ for all *m*. A raw transition frequency matrix *F* ∈ ℝ^6×6^ was then populated via a trial-wise moving count:

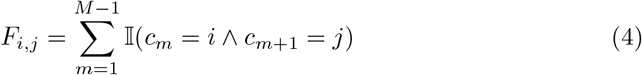

where I(·) represents an indicator function evaluating to 1 if the condition is met and 0 otherwise. Individual differences in cumulative replay frequency were controlled for by normalizing *F* into a transition matrix *T* .

Replay template matching was subsequently performed by calculating the cosine similarity (*d*_cos_) between the empirical transition matrix (*T* ) and three predefined theoretical structural templates: obvious-rule (*O*), hidden-rule (*H*), and random reactivation (*R*). These reference matrices were constructed from adjacent state pairs defined by the task rules. Empirical evidence indicates that memory replay can manifest in both forward and backward directions [12, 23]; therefore, these structures were organized as bidirectional transitions by summing their directional forward (*fwd*) and backward (*bwd* = *fwd*^*T*^ ) adjacency profiles before normalization:

1. **Obvious-rule Template (***O*): Formulated from the underlying deterministic graph structure of the task, capturing the step-by-step sequential transition patterns.

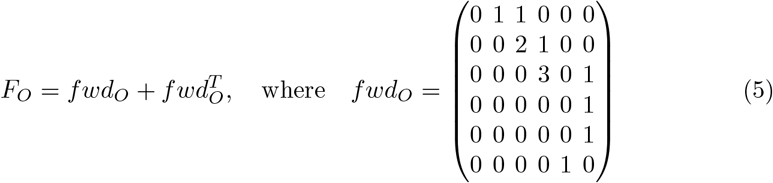
2. **Hidden-rule Template (***H*): Modeled to capture the hidden structural shortcut discovered by participants in the insight group.

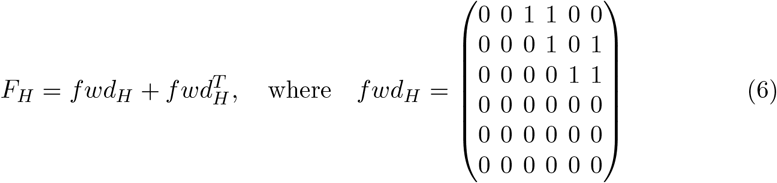
3. **Random Template (***R*): Modeled as an all-to-all connectivity profile representing uniform, unstructured transitions across task items. Self-transitions were excluded.

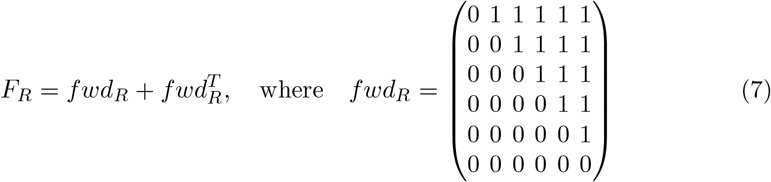

Each base frequency matrix (*F*_*O*_, *F*_*H*_, *F*_*R*_) was subsequently normalized. The resulting cosine metrics—*d*_cos_(*T, O*), *d*_cos_(*T, H*), and *d*_cos_(*T, R*)—quantified the degree of structural alignment with each rule type, serving as the predictors for subsequent insight decoding models.

#### 4.5.4 Insight Prediction

Participant-level transition features were used to train binary classifiers distinguishing insight from no-insight participants to determine whether replay dynamics could predict subsequent insight. Feature vectors comprised cosine similarity scores derived from the template-matching analysis (*d*_cos_(*T, O*), *d*_cos_(*T, H*), and *d*_cos_(*T, R*)).

Insight prediction was performed independently for each temporal replay lag condition using a k-nearest neighbors classifier (*k* = 4) with inverse-distance weighting and the Euclidean distance metric, as implemented in `scikit-learn` (v1.8.0) [55]. The design matrix comprised the replay template-matching features together with all corresponding interaction terms.

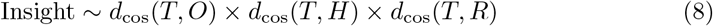

Permutation-based feature importance analysis was subsequently conducted to quantify the relative contribution of each replay attribute to overall classification performance. For each independent cross-validation fold, baseline PR-AUC was first established on the held-out validation set. Individual feature vectors were then randomly shuffled across participants while preserving the structural integrity of all remaining features, allowing the corresponding reduction in performance to be explicitly isolated:

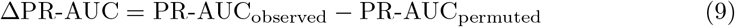

As this feature permutation was applied upstream of interaction-term generation within the computational pipeline, shuffling a given baseline replay feature simultaneously disrupted all associated two-way and three-way interaction terms involving that specific metric. Consequently, the resulting importance estimates capture the unified contribution of both the main and interaction effects. This procedure was repeated 20 times per feature and averaged across all cross-validation folds, where larger values of ΔPR-AUC greater functional importance to the predictive model.

Characterization of the relationship between the latent transition features and decoder outcomes was conducted via post-hoc ordinary least squares (OLS) regression using the mean out-of-fold (OOF) predicted probabilities as the dependent variable:

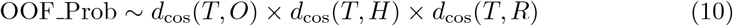

This secondary parametric model was constructed exclusively for feature interpretation, given that the internal decision architecture of the primary non-parametric *k*-nearest neighbors classifier cannot be directly evaluated through structural weights. Heteroscedasticity-consistent (HC3) standard errors were applied during parameter estimation to ensure robust statistical inference against non-constant error variance, while potential multicollinearity was monitored by calculating variance inflation factors (VIF) for each independent term.

### 4.6 Source-space connectivity analysis

An empirical noise covariance matrix was computed from the pre-rest session data. Cortical source reconstruction was performed on the standard `fsaverage` template brain via a three-layer boundary element model (BEM) and an `ico-5` source space layout [56]. The corresponding forward solution was computed with a strict minimum source-to-skull distance constraint of 5 mm to minimize volume conduction artifacts.

Distributed source estimates were obtained using exact low-resolution electromagnetic tomography (eLORETA) [57], with source orientations fully constrained to the cortical surface normal. Source-level activity was subsequently parcellated into anatomically defined cortical regions using the 64-region Desikan–Killiany atlas [58]. Functional connectivity between these parcellated cortical nodes was quantified via the weighted phase lag index (wPLI) [21], implemented within the `mne-connectivity` (v0.7.0) software library [51, 52].

Weighted connectivity matrices were calculated independently for each participant across standard canonical frequency bands: theta (4–8 Hz), alpha (8–13 Hz), beta (13– 30 Hz), low-gamma (30–50 Hz), and high-gamma (50–100 Hz). These matrices were subsequently reduced to a minimum spanning tree (MST) graph topology, systematically eliminating arbitrary thresholding-related biases and density variations during network analysis [59].

### 4.7 Statistical analysis

Normality of the data distribution was assessed using the Shapiro–Wilk test prior to any statistical comparisons. Statistical tests were subsequently chosen based on whether these underlying distributional assumptions of normality were satisfied.Statistical differences in localizer model decoding accuracy were evaluated using a linear mixed-effects model (LMM) implemented in `statsmodels` (v0.14.0) [60]:

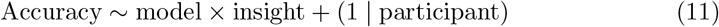

Post-hoc pairwise comparisons between models were performed using two-tailed Wilcoxon signed-rank tests, with *p*-values adjusted using the Bonferroni correction.

Insight prediction performance was quantified using the area under the precision– recall curve (PR-AUC) across 10 repetitions of stratified 10-fold cross-validation. Statistical significance was assessed using permutation testing with 10,000 random label shuffles. For lag conditions exhibiting significant predictive performance, permutation-based feature importance was computed as the reduction in PR-AUC following random shuffling of individual feature vectors.

Group-level topological differences in source-space functional connectivity were evaluated using the network-based statistic (NBS) analysis [61]. The MST-reduced connectivity matrices from the insight and no-insight groups were compared independently within each frequency band via two-tailed unpaired *t*-tests. Selection of the primary edge-level test-statistic threshold was informed by the expected effect size (*d* = 0.65) and adjusted dynamically relative to group sample sizes. Specifically, this operational threshold was set at *t*_*threshold*_ = 2.353, representing the larger value between the expected *t*-statistic derived from Cohen’s *d* and the critical student’s *t*-value corresponding to *α* = 0.05. Family-wise error rate control at the subnetwork level was subsequently enforced via non-parametric permutation testing using 10,000 random permutations.

## Supplementary information

## Acknowledgements

The authors would like to thank Dr. Miyoko Street, Megumi Inoue, and Minako Inoue for their assistance with data collection. The authors also thank members of the Chao Lab for their valuable feedback and comments. The authors further acknowledge the contribution of all study participants and the support of the participant recruitment service provided by https://www.jikken-baito.com. The authors acknowledge the use and adaptation of illustrations created by Takashi Mifune and distributed through Irasutoya (https://www.irasutoya.com/)in the experimental materials.

## Declarations

- Funding This work was supported by the World Premier International Research Center Initiative (WPI), MEXT, Japan (to Z.C.C.) and the IRCN-Daikin SCP (to Z.C.C.). The funders had no role in study design, data collection, and analysis, the decision to publish, or the preparation of the manuscript.
- Conflict of interest/Competing interests (check journal-specific guidelines for which heading to use) The authors declare no competing interests.
- Ethics approval and consent to participate All participants provided written informed consent prior to the experiment. The protocol was approved by the Ethics Committee of the University of Tokyo (IRB Number: E2025ALS288) and conducted in accordance with the Declaration of Helsinki.
- Author contribution C.P. (conceptualization, data curation, formal analysis, investigation, methodology, software, validation, writing–original draft, writing–review and editing) and Z.C. (conceptualization, project administration, resources, methodology, validation, supervision, writing–review and editing).

