## Supplementary for "From Subconscious to Insight: Decoding the Incubation Process in Creative Problem Solving"

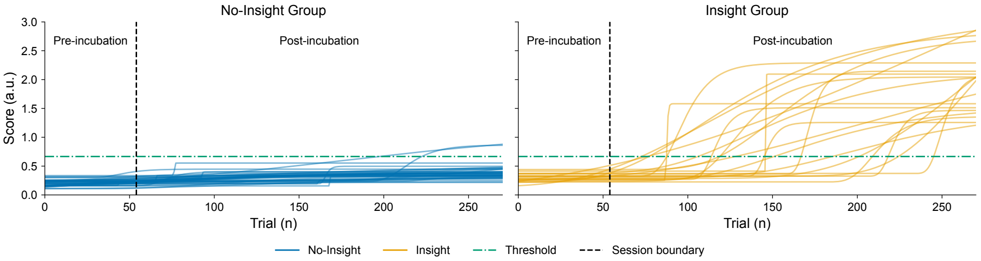


**Supplementary Fig. 1**: **Gompertz fitted score.** Each line represents a single participant in the insight (blue solid lines) and no-insight (orange solid lines) groups. The vertical black dashed line separates the pre-incubation and post-incubation sessions.

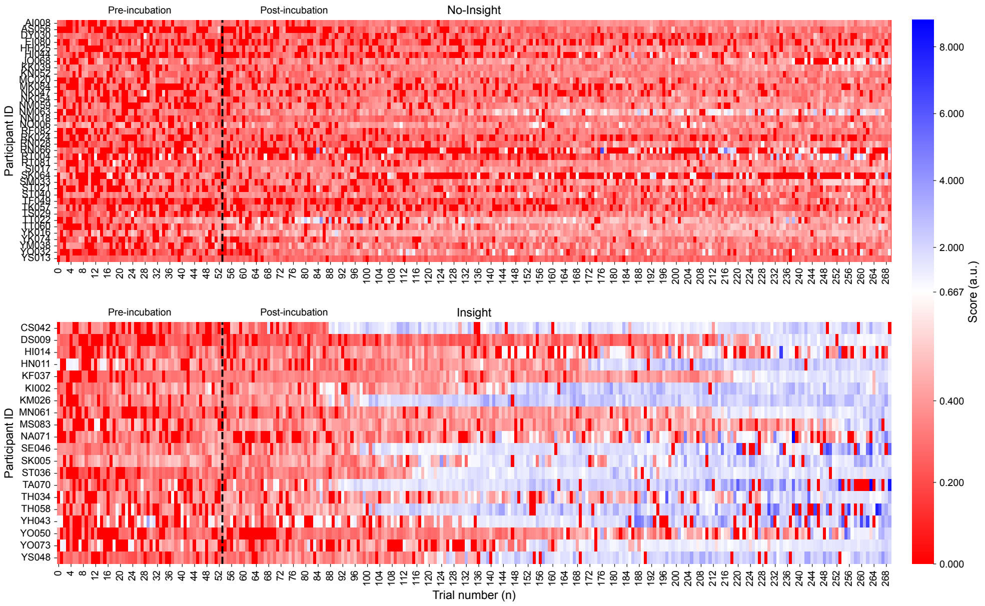


**Supplementary Fig. 2**: **Raw behavioral scores across participants.** Heatmaps show behavioral scores for each participant across trials. Red indicates scores below the threshold of 1/1.5, whereas blue indicates scores above this threshold. The vertical dashed line marks the boundary between the pre-incubation and post-incubation sessions. The top panel corresponds to the no-insight group, and the bottom panel corresponds to the insight group.

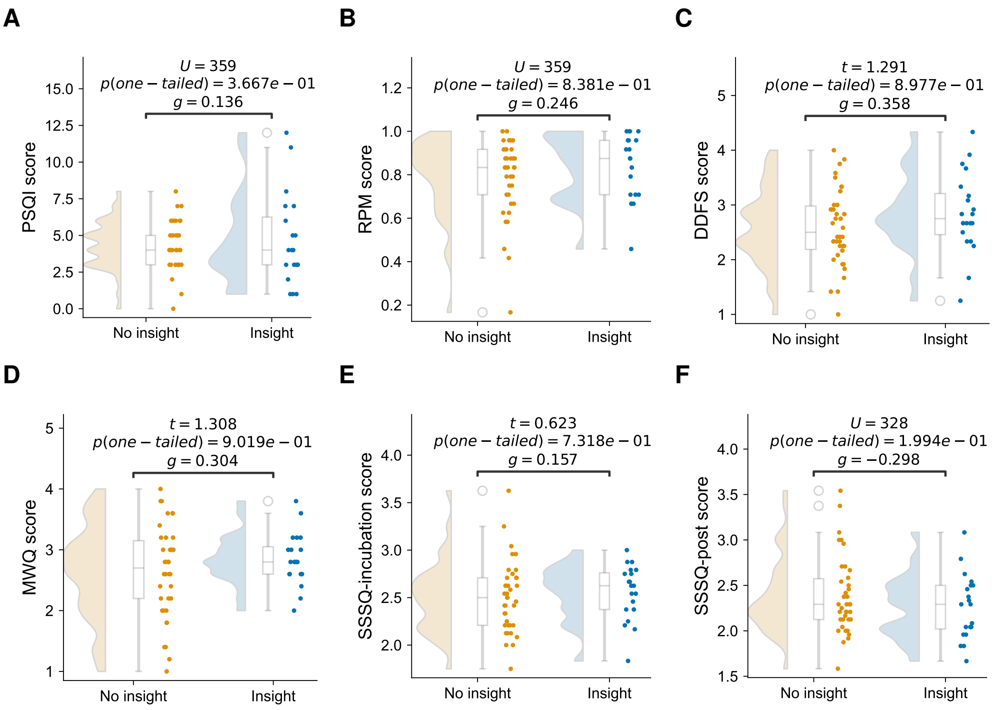


**Supplementary Fig. 3**: **Demographic metrics and baseline score distributions across behavioral groups.** There are no significant between-group differences across any of the evaluated psychometric or state parameters. **A** PSQI distributions. **B** RPM results; note that for the RPM metric, only the number of correct answers was used to calculate the final score. **C** DDFS profiles. **D** MWQ ratings. **E** SSSQ values recorded after completing the pre-incubation session. **F** SSSQ values recorded after completing the post-incubation session. Exact statistical values for group comparisons—including the significance level ($p$), Mann–Whitney test statistic ($U$), Student's $t$-statistic ($t$) and Hedges' $g$ effect size ($g$)—are displayed directly at the top of each corresponding subpanel for direct comparison.

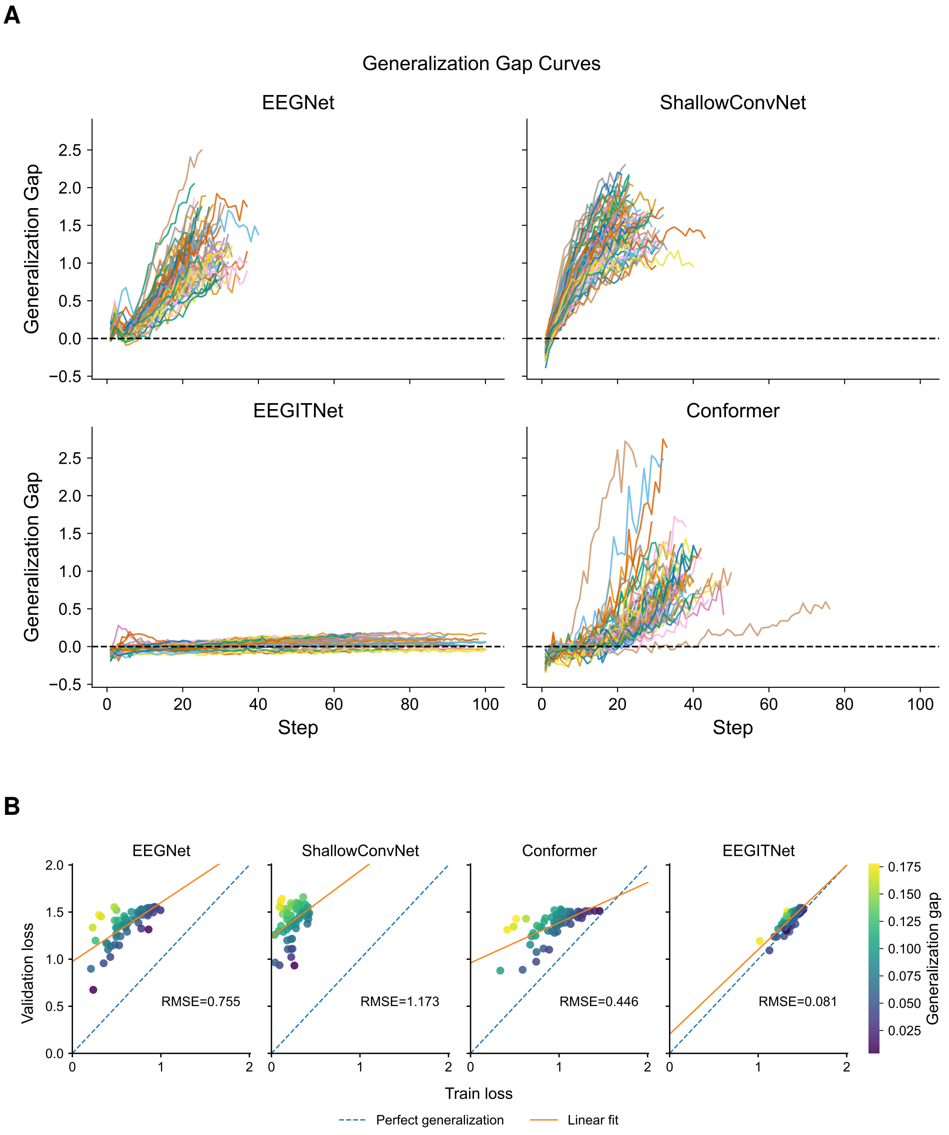


**Supplementary Fig. 4**: **Generalization performance across models. A** Generalization gap for each subject, computed as the difference between validation and training loss across optimization steps. **B** Generalization performance across all models, assessed by the relationship between training and validation loss.

**Supplementary Table 1**: EEG Data Augmentation Pipeline Methods, Hyperparameters, and Functional Mechanisms

| Augmentation Method | Hyperparameters | Functional Mechanism |
|---|---|---|
| **SegmentationReconstruction** | $p = 0.1$, $N_{\text{segments}} = \max\left(\{f \in \mathbb{Z} : f \mid N_{\text{samples}}, f \leq \sqrt{N_{\text{samples}}}\}\right)$ | Divides the continuous EEG signal into localized temporal windows, shuffles their order, and smoothly reconstructs the boundary segments. |
| **FrequencyShift** | $p = 0.1$, $\Delta f_{\max} = 2$ Hz, $f_s = \text{sfreq}$ | Transports the signal into the frequency domain and rigidly shifts all spectral components up or down by a random delta up to 2 Hz. |
| **FTSurrogate** | $p = 0.1$, phase_noise_mag = 1.0, channel_indep = False | Performs a Fast Fourier Transform (FFT), randomizes the phase spectrum while keeping the original amplitude spectrum intact, and transforms back into the time domain. |
| **ChannelsDropout** | $p = 0.1$ | Completely zeroes out / deactivates a random subset of spatial recording channels for the duration of the trial to force spatial invariance. |
| **SmoothTimeMask** | $p = 0.1$, mask_len = 50 samples | Replaces a contiguous block of 50 temporal samples with zeros, using a smooth window fade to avoid generating artificial, sharp high-frequency edge artifacts. |
| **TimeReverse** | $p = 0.1$ | Inverts the sequence chronologically along the time axis (flips the trial backwards from end to start). |
| **GaussianNoise** | $p = 0.1$, $\sigma = 0.1$ | Injects stochastic additive white Gaussian noise drawn from a distribution with a standard deviation ($\sigma$) of 0.1 to simulate ambient sensor noise. |

**Supplementary Table 2**: Linear mixed-effects model predicting decoding accuracy as a function of model architecture and insight group.

| **Fixed Effects** | **Estimate** | **SE** | $z$ | $p$ | **95% CI** | |
|---|---|---|---|---|---|---|
| Intercept | 0.509 | 0.015 | 34.55 | 1.578e-261 | 0.480 | 0.538 |
| Model | -0.029 | 0.004 | -7.90 | 2.803e-15 | -0.037 | -0.022 |
| Insight Group | -0.007 | 0.025 | -0.28 | 0.783 | -0.056 | 0.042 |
| Model : Insight Group | 0.005 | 0.006 | 0.71 | 0.475 | -0.008 | 0.017 |
| **Random Effects** | | | | | | |
| Participant variance ($\sigma^2_{\text{participant}}$) | 0.006 | 0.030 | — | — | — | — |
| Residual variance ($\sigma^2_{\text{residual}}$) | 0.0026 | — | — | — | — | — |
| **Model Information** | | | | | | |
| Dependent variable | *Accuracy* | Observations: 232 | | | Participants: 58 | |
| Estimation | REML | Log-likelihood: 277.412 | | | Group size: 4 / 4 / 4.0 | |

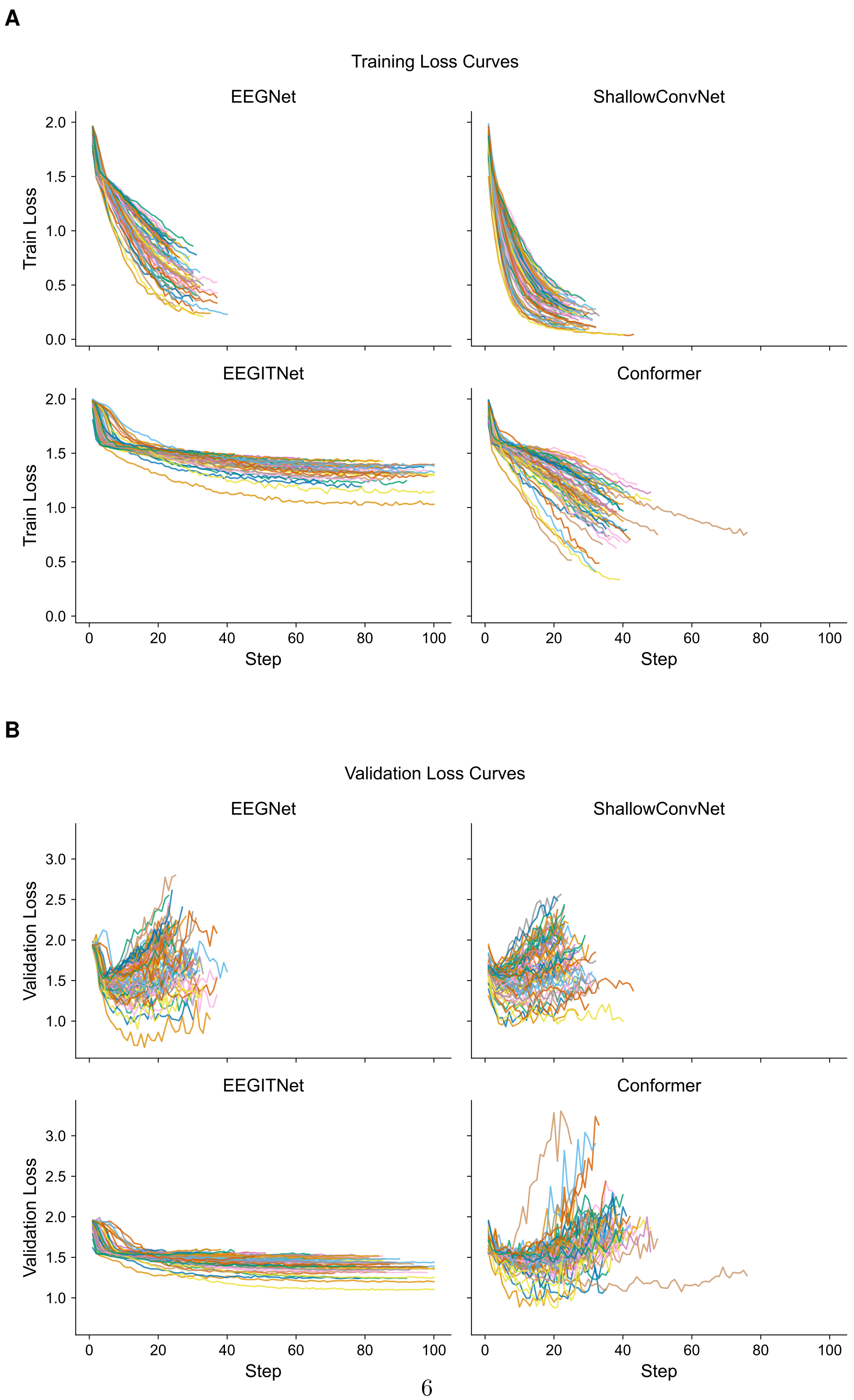




**Supplementary Fig. 5**: **Training and validation loss dynamics across subjects and models. A** Training loss curves over optimization steps for each subject, separated by model architecture. **B** Validation loss curves over optimization steps for each subject, separated by model architecture.

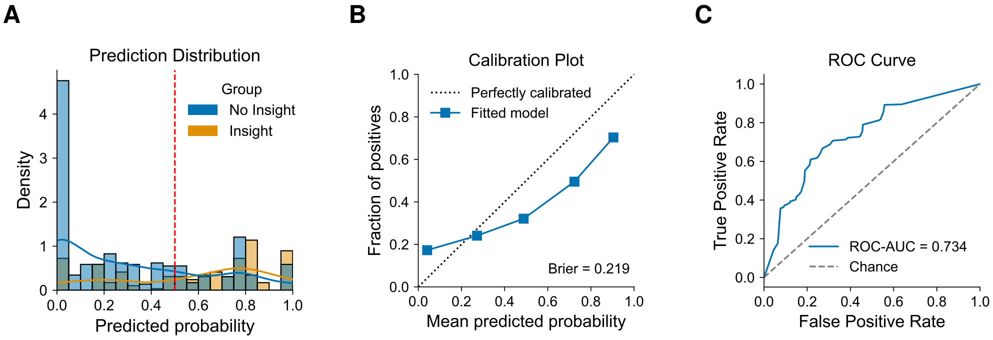


**Supplementary Fig. 6**: **Predicted probability distributions and model calibration for predicting subsequent insight from memory replay features.** **A** Distribution of predicted probabilities for insight and no-insight participants, illustrating separation between groups. **B** Calibration curve showing the relationship between mean predicted probabilities and the observed fraction of positive outcomes. **C** Receiver operating characteristic (ROC) curve for the model, with area under the curve (AUC) indicating discriminative performance ($p = 7.900e - 3$).

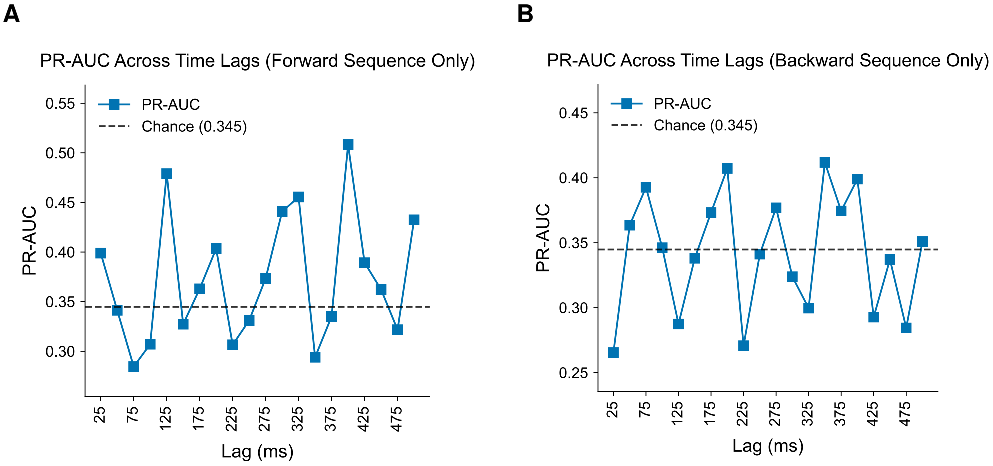


**Supplementary Fig. 7**: **Area under the precision-recall curve across temporal lags.** Both **A** forward sequence and **B** backward sequence decoding performance show no significant deviations from the empirically derived chance baseline.

**Supplementary Table 3**: Ordinary least squares regression predicting out-of-fold prediction scores from cosine-distance features and their interaction terms. Robust HC3 standard errors were used.

| **Predictor** | **Estimate** | **SE** | $z$ | $p$ | **95% CI** | |
|---|---|---|---|---|---|---|
| Intercept | 0.478 | 0.044 | 10.796 | 3.583e-27 | 0.391 | 0.564 |
| **Main Effects** | | | | | | |
| $d_{\cos}(T, O)$ | 0.185 | 0.416 | 0.444 | 0.657 | -0.630 | 0.999 |
| $d_{\cos}(T, H)$ | 1.448 | 0.430 | 3.372 | 7.469e-4 | 0.606 | 2.290 |
| $d_{\cos}(T, R)$ | -0.114 | 0.402 | -0.283 | 0.777 | -0.902 | 0.674 |
| **Two-Way Interactions** | | | | | | |
| $d_{\cos}(T, O)$ : $d_{\cos}(T, H)$ | 0.746 | 2.712 | 0.275 | 0.783 | -4.571 | 6.062 |
| $d_{\cos}(T, O)$ : $d_{\cos}(T, R)$ | -8.578 | 2.729 | -3.143 | 1.673e-3 | -13.927 | -3.228 |
| $d_{\cos}(T, H)$ : $d_{\cos}(T, R)$ | -2.195 | 2.765 | -0.794 | 0.427 | -7.615 | 3.225 |
| **Three-Way Interaction** | | | | | | |
| $d_{\cos}(T, O)$ : $d_{\cos}(T, H)$ : $d_{\cos}(T, R)$ | -26.876 | 5.800 | -4.634 | 3.586e-6 | -38.244 | -15.509 |
| **Model Statistics** | | | | | | |
| Dependent variable | *OOF_Prob* | $N = 58$ | | | $R^2 = 0.412$ | |
| Adjusted $R^2$ | 0.330 | $F = 9.478$ | | | $p = 1.989e - 7$ | |
| Log-likelihood | -2.172 | AIC = 20.34 | | | BIC = 36.83 | |
| Covariance type | HC3 | Durbin–Watson = 2.007 | | | Condition No. = 212 | |

**Supplementary Table 4**: EEGNet Model Architecture and Layer Specifications [1]

| Layer / Operation (Type) | Input Shape | Output Shape | Kernel Shape | Param # |
|---|---|---|---|---|
| EEGNet (Input) | [1, 64, 251] | [1, 7] | – | – |
| ↪ Ensure4d | [1, 64, 251] | [1, 64, 251, 1] | – | – |
| ↪ Rearrange | [1, 64, 251, 1] | [1, 1, 64, 251] | – | – |
| ↪ Conv2d | [1, 1, 64, 251] | [1, 64, 64, 252] | [1, 64] | 4,096 |
| ↪ BatchNorm2d | [1, 64, 64, 252] | [1, 64, 64, 252] | – | 128 |
| ↪ Depthwise Conv2d (Constraint) | [1, 64, 64, 252] | [1, 128, 1, 252] | [64, 1] | 8,192 |
| ↪ BatchNorm2d | [1, 128, 1, 252] | [1, 128, 1, 252] | – | 256 |
| ↪ ELU | [1, 128, 1, 252] | [1, 128, 1, 252] | – | – |
| ↪ AvgPool2d | [1, 128, 1, 252] | [1, 128, 1, 63] | [1, 4] | – |
| ↪ Dropout | [1, 128, 1, 63] | [1, 128, 1, 63] | – | – |
| ↪ Separable Conv2d | [1, 128, 1, 63] | [1, 128, 1, 64] | [1, 16] | 2,048 |
| ↪ Pointwise Conv2d | [1, 128, 1, 64] | [1, 128, 1, 64] | [1, 1] | 16,384 |
| ↪ BatchNorm2d | [1, 128, 1, 64] | [1, 128, 1, 64] | – | 256 |
| ↪ ELU | [1, 128, 1, 64] | [1, 128, 1, 64] | – | – |
| ↪ AvgPool2d | [1, 128, 1, 64] | [1, 128, 1, 8] | [1, 8] | – |
| ↪ Dropout | [1, 128, 1, 8] | [1, 128, 1, 8] | – | – |
| ↪ Classification Head (Sequential) | [1, 128, 1, 8] | [1, 7] | – | – |
| ↪ ↪ Conv2d | [1, 128, 1, 8] | [1, 7, 1, 1] | [1, 8] | 7,175 |
| ↪ ↪ Output Formatting (Squeeze) | [1, 7, 1, 1] | [1, 7] | – | – |

**Model Computational Complexity Statistics:**

| | |
|---|---|
| Total / Trainable Parameters: 38,535 | Total Mult-Adds: 67.25 MB |
| Input Size: 0.06 MB | Forward/Backward Pass Size: 16.97 MB |
| Parameter Memory Footprint: 0.12 MB | **Estimated Total Size:** 17.16 MB |

**Supplementary Table 5**: ShallowConvNet Model Architecture and Layer Specifications [2]

| Layer / Operation (Type) | Input Shape | Output Shape | Kernel Shape | Param # |
|---|---|---|---|---|
| ShallowConvNet (Input) | [1, 64, 251] | [1, 7] | – | – |
| ↪ Ensure4d | [1, 64, 251] | [1, 64, 251, 1] | – | – |
| ↪ Rearrange | [1, 64, 251, 1] | [1, 1, 251, 64] | – | – |
| ↪ CombinedConv | [1, 1, 251, 64] | [1, 40, 227, 1] | – | 103,440 |
| ↪ BatchNorm2d | [1, 40, 227, 1] | [1, 40, 227, 1] | – | 80 |
| ↪ Expression (Square Activation) | [1, 40, 227, 1] | [1, 40, 227, 1] | – | – |
| ↪ AvgPool2d | [1, 40, 227, 1] | [1, 40, 11, 1] | [75, 1] | – |
| ↪ SafeLog (Log Activation) | [1, 40, 11, 1] | [1, 40, 11, 1] | – | – |
| ↪ Dropout | [1, 40, 11, 1] | [1, 40, 11, 1] | – | – |
| ↪ Classification Head (Sequential) | [1, 40, 11, 1] | [1, 7] | – | – |
| ↪ ↪ Conv2d | [1, 40, 11, 1] | [1, 7, 1, 1] | [11, 1] | 3,087 |
| ↪ ↪ Output Formatting (Squeeze) | [1, 7, 1, 1] | [1, 7] | – | – |

**Model Computational Complexity Statistics:**

| | |
|---|---|
| Total / Trainable Parameters: 106,607 | Total Mult-Adds: 0.00 MB |
| Input Size: 0.06 MB | Forward/Backward Pass Size: 0.07 MB |
| Parameter Memory Footprint: 0.01 MB | **Estimated Total Size:** 0.15 MB |

**Supplementary Table 6**: EEGConformer Model Architecture and Layer Specifications [3]

| Layer / Operation (Type) | Input Shape | Output Shape | Kernel Shape | Param # |
|---|---|---|---|---|
| EEGConformer (Input) | $[1, 64, 251]$ | $[1, 7]$ | – | – |
| **1. Patch Embedding Module** | | | | |
| ↪ Conv2d (Temporal Filtering) | $[1, 1, 64, 251]$ | $[1, 40, 64, 227]$ | $[1, 25]$ | 1,040 |
| ↪ Conv2d (Spatial Filtering) | $[1, 40, 64, 227]$ | $[1, 40, 1, 227]$ | $[64, 1]$ | 102,440 |
| ↪ BatchNorm2d + ELU | $[1, 40, 1, 227]$ | $[1, 40, 1, 227]$ | – | 80 |
| ↪ AvgPool2d | $[1, 40, 1, 227]$ | $[1, 40, 1, 11]$ | $[1, 75]$ | – |
| ↪ Conv2d (Projection Head) | $[1, 40, 1, 11]$ | $[1, 40, 1, 11]$ | $[1, 1]$ | 1,640 |
| ↪ Rearrange (Sequence Formatting) | $[1, 40, 1, 11]$ | $[1, 11, 40]$ | – | – |
| **2. Transformer Encoder Module** (6× **Blocks**) | | | | |
| ↪ Multi-Head Self-Attention + Residual Add | $[1, 11, 40]$ | $[1, 11, 40]$ | – | $6 \times 6,640$ |
| ↪ Feed-Forward Network + Residual Add | $[1, 11, 40]$ | $[1, 11, 40]$ | – | $6 \times 13,080$ |
| **3. Fully Connected Classification Head** | | | | |
| ↪ Linear (Flattened Sequence to Hidden) | $[1, 440]$ | $[1, 256]$ | – | 112,896 |
| ↪ ELU + Dropout | $[1, 256]$ | $[1, 256]$ | – | – |
| ↪ Linear (Hidden to Latent) | $[1, 256]$ | $[1, 32]$ | – | 8,224 |
| ↪ ELU + Dropout | $[1, 32]$ | $[1, 32]$ | – | – |
| ↪ Linear (Output Projection) | $[1, 32]$ | $[1, 7]$ | – | 231 |
| **Model Computational Complexity Statistics:** | | | | |
| Total / Trainable Parameters: 344,871 | | Total Mult-Adds: 38.62 MB | | |
| Input Size: 0.06 MB | | Forward/Backward Pass Size: 5.03 MB | | |
| Parameter Memory Footprint: 1.38 MB | | **Estimated Total Size:** 6.48 MB | | |

**Supplementary Table 7**: EEGITNet Model Architecture and Layer Specifications [4]

| Layer / Operation (Type) | Input Shape | Output Shape | Kernel Shape | Param # |
|---|---|---|---|---|
| EEGITNet (Input) | $[1, 64, 251]$ | $[1, 7]$ | – | – |
| **1. Input Preprocessing** | | | | |
| ↪ Ensure4d + Rearrange | $[1, 64, 251]$ | $[1, 1, 64, 251]$ | – | – |
| **2. Inception Feature Extraction Block** | | | | |
| ↪ Stream 1 (Sequential) | $[1, 1, 64, 251]$ | $[1, 2, 1, 251]$ | – | 168 |
| ↪ Stream 2 (Sequential) | $[1, 1, 64, 251]$ | $[1, 4, 1, 251]$ | – | 400 |
| ↪ Stream 3 (Sequential) | $[1, 1, 64, 251]$ | $[1, 8, 1, 251]$ | – | 608 |
| ↪ Average Pooling + Dropout | $[1, 14, 1, 251]$ | $[1, 14, 1, 62]$ | $[1, 4]$ | – |
| **3. Temporal Convolutional Processing Modules (_TCBlocks)** | | | | |
| ↪ _TCBlock 1 (Dilation Rate = 1) | $[1, 14, 1, 62]$ | $[1, 14, 1, 62]$ | $2 \times [1, 4]$ | $56 + 28$ |
| ↪ _TCBlock 2 (Dilation Rate = 2) | $[1, 14, 1, 62]$ | $[1, 14, 1, 62]$ | $2 \times [1, 4]$ | $56 + 28$ |
| ↪ _TCBlock 3 (Dilation Rate = 4) | $[1, 14, 1, 62]$ | $[1, 14, 1, 62]$ | $2 \times [1, 4]$ | $56 + 28$ |
| ↪ _TCBlock 4 (Dilation Rate = 8) | $[1, 14, 1, 62]$ | $[1, 14, 1, 62]$ | $2 \times [1, 4]$ | $56 + 28$ |
| **4. Dimensionality Reduction & Classification Head** | | | | |
| ↪ Conv2d Feature Projection | $[1, 14, 1, 62]$ | $[1, 28, 1, 62]$ | $[1, 1]$ | 420 |
| ↪ BatchNorm2d + ELU | $[1, 28, 1, 62]$ | $[1, 28, 1, 62]$ | – | 56 |
| ↪ AvgPool2d + Dropout | $[1, 28, 1, 62]$ | $[1, 28, 1, 15]$ | $[1, 4]$ | – |
| ↪ Flatten | $[1, 28, 1, 15]$ | $[1, 420]$ | – | – |
| ↪ Linear Classification Layer | $[1, 420]$ | $[1, 7]$ | – | 2,947 |
| **Model Computational Complexity Statistics:** | | | | |
| Total / Trainable Parameters: 5,271 | | Total Mult-Adds: 3.88 MB | | |
| Input Size: 0.06 MB | | Forward/Backward Pass Size: 3.79 MB | | |
| Parameter Memory Footprint: 0.02 MB | | **Estimated Total Size:** 3.88 MB | | |
